# Increase in ribosomal proteins activity: Translational reprogramming in *Vanilla planifolia* Jacks., against *Fusarium* infection

**DOI:** 10.1101/660860

**Authors:** Marco Tulio Solano de la Cruz, Jacel Adame-García, Josefat Gregorio-Jorge, Verónica Jiménez-Jacinto, Leticia Vega-Alvarado, Lourdes Iglesias-Andreu, Esteban Elías Escobar-Hernández, Mauricio Luna-Rodríguez

## Abstract

**Background:** Upon exposure to unfavorable environmental conditions, plants need to respond quickly to maintain their homeostasis. For instance, physiological, biochemical and transcriptomical changes must occur during interactions with pathogens, this causing the triggering of pathogen- and plant-derived molecules. In the case of *Vanilla planifolia* Jacks., a worldwide economically important crop, it is susceptible to *Fusarium oxysporum* f. sp. *vanillae*. This pathogen causes root and stem rot in vanilla plants that finally leads to plant death. To further investigate how vanilla plants respond at the transcriptional level upon infection with *F.oxysporum* f. sp. *vanillae*, we employed the RNA-Seq approach to analyze the dynamics of whole-transcriptome changes during two-time frames of the infection.

**Results:** Analysis of global gene expression profiles indicated that a major transcriptional change occurs at 2 dpi, in comparison to 10 dpi, whereas 3420 genes were found with a differential expression at 2 dpi, only 839 were identified at 10 dpi. The analysis of the transcriptional profile at 2 dpi suggests that vanilla plants prepare to counter the infection by gathering a pool of translational regulation-related transcripts.

**Conclusions:** We propose that the plant-pathogen interaction at early stages causes a transcriptional reprogramming coupled with a translational regulation. Altogether, this study provides the identification of molecular players that could help to fight the most damaging disease of vanilla, where ribosomal proteins and regulation of the translational mechanism are critical. These are insights into the defense responses of *V. planifolia* Jacks., providing the basis for the understanding of the plant early response towards biotic stress.

## INTRODUCTION

Throughout evolution, plants have developed multiple defense strategies to cope with pathogens. The first defense line consists of pre-existing physical and chemical barriers, which restrict their entry. In addition to these constitutive barriers, plants have developed an immune response mechanism that is based on the detection of elicitor compounds derived from pathogens, known as Pathogen-Associated Molecular Patterns (PAMPs) (Shiu et al., 2004). Such defense response activated by the PAMPs or PAMP-Triggered Immunity (PTI), usually restricts the proliferation of the pathogen (Zhang and Zhou, 2010; Bigeard et al, 2015; Ausubel, 2005; Boller and Felix, 2009; Yamaguchi and Huffaker, 2010). However, some pathogens have circumvented this response by developing effector proteins that interfere or suppress PTI (Macho and Zipfel, 2015; Guo et al., 2009; Zipfel, 2014). In this sense, the so-called co-evolutionary ‘arms race’ between plants and pathogens has defined the establishment of the Effector-Triggered Immunity (ETI), a defense line that begins with the recognition of PAMPs by plant pattern recognition receptors (PRRs) (Jones and Dangl, 2006). The signals generated by PRRs are transduced through Mitogen-activated Protein Kinases (MAPKs), which in turn activate transcription factors for gene regulation that leads to a proper plant defense response. Among the plant responses, the Hypersensitive Response (RH), the programmed cell death, the expression of proteins related to pathogenesis or the lignification of the cell wall are included (Chen and Ronald, 2011; Chiang and Coaker, 2015; Spoel and Dong, 2012; Naito et al., 2007; Clough et al., 2000; Coll et al., 2010).

*Vanilla planifolia* Jacks., is one of the most economically relevant orchids. It is produced extensively in several countries and is the main natural source of one of the most widely used flavoring agents in the world, vanillin (Anilkumar, 2004; Hernández-Hernández, 2011). Its cultivation has spread throughout the world, with Madagascar and Indonesia as the leaders of annual production (35.5% and 34.5%, respectively), followed by China (13.7%) and Papua New Guinea (4.1%) (De La Cruz Medina et al., 2009; Kalimuthu et al., 2007; Dignum et al. 2001; Roling et al. 2001; Pinaria and Burges, 2010). Although Mexico is the center of domestication and diversification of this crop, vanillin production is positioned in the fifth place, contributing to only 4.0% of world production (Hernández-Hernández, 2011). Importantly, vanilla plants are susceptible to parasites and pathogens. The most lethal pathogen that afflicts vanilla is *Fusarium oxysporum* f. sp. *vanillae*, a pathogenic form of the genus *Fusarium* that specifically infects this plant (Ramírez-Mosqueda et al, 2018; Kalimuthu et al.,2006; Pinaria and Burges, 2010). This pathogen causes root and stem rot, as well as the colonization of vascular tissues that finally leads to plant death. Several studies indicate that *V. planifolia* Jacks., has a high susceptibility and incidence of *Fusarium oxysporum* f. sp. *vanillae* (Bhai and Dhanesh, 2008, Pinaria and Burges, 2010, Fravel et al., 2003). For instance, infection of vanilla plants by this pathogen is capable of destroying 65% of the plantation (Ramírez-Mosqueda et al, 2018; Kalimuthu et al., 2006; Pinaria and Burges, 2010). The lack of genetic variability of *V. planifolia* Jacks is another factor that worsens the scenario (Bory et al., 2008; Lubinsky et al., 2008; Ramírez-Mosqueda et al, 2018). Thus, given the economic importance of *V. planifolia* Jacks., in the last decade several efforts have tried to elucidate the overall plant response upon infection by this pathogen (Bai et al, 2013; Li et al., 2012; Xing et al., 2016). Moreover, since inferences from mRNA expression data are valuable as it reflects changes with a biological meaning, we looked into the transcriptome of *V. planifolia* roots exposed to *Fusarium oxysporum* f. sp. *vanillae*, to figure out the responsive mechanisms at early (2 days after inoculation, 2dpi) and later (10 days after inoculation, 10 dpi) stages of infection. Gene expression profiles indicated that major transcriptional changes occurs at 2 dpi. Accordingly, vanilla plants prepare to counter the infection by gathering a pool of translational regulation-related transcripts. Altogether, this study provides the identification of molecular players that could help to fight the most damaging disease of vanilla.

## RESULTS

### Assembly of the transcriptome of *V. planifolia* roots exposed to *F. oxysporum* f. sp. *vanillae*

The transcriptome of vanilla roots exposed to *F. oxysporum* f. sp. *vanillae* was assessed with Illumina sequencing at 2 and 10 dpi. A total of 12 cDNA libraries were paired-end sequenced using the NextSeq 500 system. Sequencing data of these libraries were obtained, corresponding to three biological replicates for each control and treatment used, as well as the two frames of time evaluated. In brief, six libraries corresponding to control and treatment at 2 dpi, as well as six libraries at 10 dpi, produced more than 204 million reads. Pre-processing of raw sequencing reads was carried out with FastQC, which indicated a good per base quality. Filtering reads that correspond to the pathogen used in the treatments at 2 and 10 dpi, discarded 5.33 and 39%, respectively. On the other hand, even that control plants were not inoculated with the fungus, 5 (2 dpi) and 6.48% (10 dpi) of reads aligned to the genome of *F. oxysporum* f. sp. *lycopersici*. Such reads were excluded from subsequent analyzes. The *de novo* transcriptome assembly of vanilla was carried out, resulting in 45,000 transcripts, associated to approximately 11,000 unigenes (Figure 1). The generated transcripts were mapped against the plant databases, using the BUSCO software, obtaining about 99% of complete orthologous. Figure 1 shows the result of the annotation of the vanilla transcriptome, with blast2go, we found annotation for about 11,000 unigenes; of the total of more than 44000. Highlighting that we find represented in the transcriptome, within the main functional categories of gene ontology obtained, a greater number of transcripts related to the development, growth, cell proliferation, signaling, response to stimuli, and the response to stress in plants. Altogether, the transcriptome of *V. planifolia* roots exposed to *F. oxysporum* f. sp. *vanillae* revealed dynamical changes in genes associated to several cellular processes.

**Figure 1.**
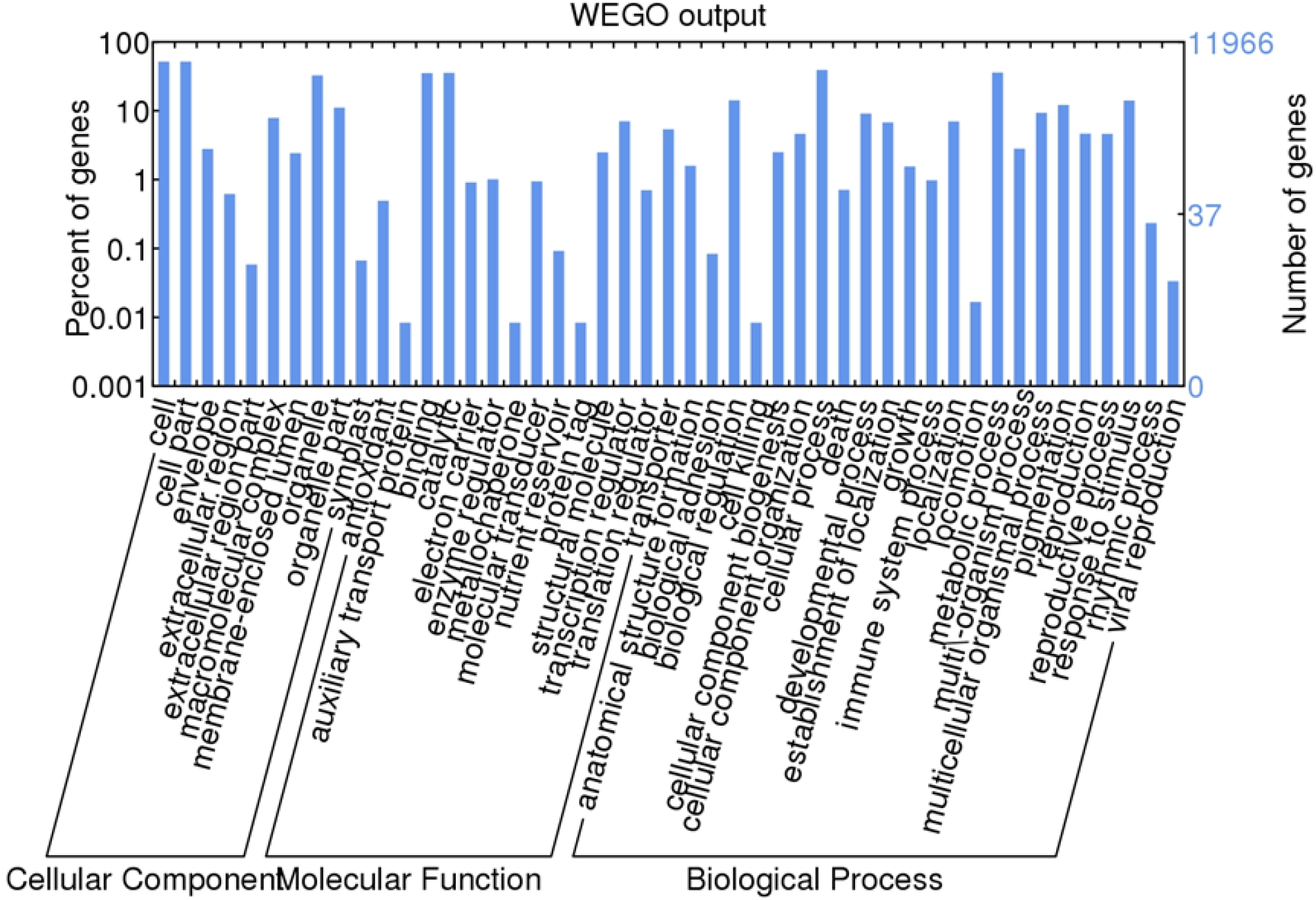
Annotation and gene ontology of the *de novo* transcriptome of *Vanilla planifolia* Jacks. Main categories of gene ontology determined by the Blast2go software, using all the annotated unigenes. The number of genes corresponding to each functional category found is observed.

### Analysis of gene expression in response to *F. oxysporum* f. sp. *vanillae* infection

For the identification of the unigenes showing changes in expression levels in the different libraries, in to both treatments (2 and 10 dpi respectively), in relation to the libraries of the control group, the differential expression analysis was carried out. As a result of the differential expression analysis performed with the DESeq, DESeq2, EdgeR and NOISeq methods. For libraries corresponding to 2 dpi, 2310 DEG genes were obtained with the DESeq method, 3420 DEG genes with the EdgeR method, while with the method NOISeq 4080 DEG genes were obtained, finally with the DESeq2 method only 1702 differentially expressed genes were obtained. These results were contrasted by a Venn diagram shown in 44. S1.

On the other hand, with regard to treatment at 10 dpi, with the DESeq method 812 DEG genes were obtained, while with the EdgeR method 881 DEG genes were obtained, similarly with the NOIseq method, 839 DEG genes were obtained, in contrast to the DESeq2 method, where only 534 DEG genes were obtained (Figure supplementary. S1). It should be noted that the unigenes differentially expressed in the libraries corresponding to the different treatments are diametrically different, since only 5 unigenes are shared between both treatments. Likewise, differentially expressed unigenes turn out to be different and specific to each treatment, and also very diverse in their function to the processes in which they participate in the plant cell (Figure 2, Figure supplementary. S2, Figure supplementary. S3, and Figure supplementary. S4).

**Figure 2.**
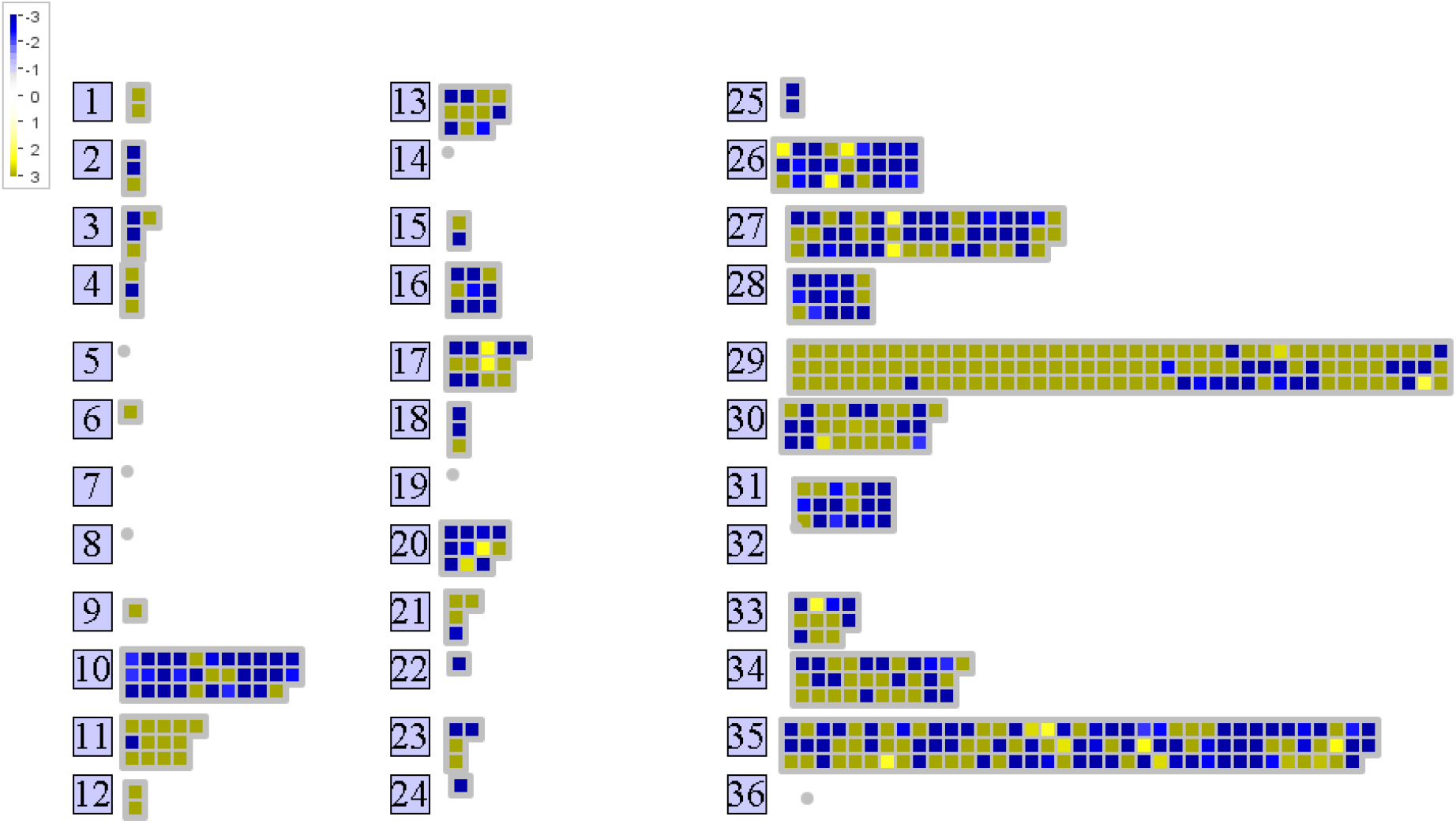
Global profiles of expression of genes differentially expressed in *Vanilla planifolia* Jacks., At 2 dpi of the infection caused by *Fusarium oxysporum* f. sp. *vanillae*. Heat map indicating the expression profiles of the annotated DEG unigenes, corresponding to the 2dpi. The numbers in the figure correspond to different categories of gene ontology, as described below: 29 protein, 35 not assigned, 11 lipid metabolism, 10 cell wall, 27 RNA, 20 stress, 25 C1-metabolism, 26 misc, 16 secondary metabolism, 28 DNA, 12 N-metabolism, 24 Biodegradation of Xenobiotics, 6 gluconeogenese/ glyoxylate cycle, 22 polyamine metabolism, 34 transport, 31 cell, 21 redox, 18 Co-factor and vitamine metabolism, 9 mitochondrial electron transport / ATP synthesis, 13 amino acid metabolism, 1 PS, 23 nucleotide metabolism, 15 metal handling, 33 development, 3 minor CHO metabolism, 17 hormone metabolism, 30 signalling, 4 glycolysis.

Based on the results of the Venn diagrams, where the DEG genes are contrasted determined with the different methods tested. We selected the EdgeR method for the subsequent analysis of differentially expressed genes in both treatments; since this method includes the great majority of the gene retrieved with the different methods. From these unigenes, determined by EdgeR, two listings of differentially expressed genes were then obtained: one corresponding to the 2dpi treatment, and the other corresponding to the 10dpi treatment. Supplemental Tables 2 and 3 show a list of the 100 genes with the highest logFC that present annotation, corresponding to treatment 2dpi. While Supplemental tables 5 and 6 show a list of the 100 genes with the highest logFC that present annotation, corresponding to treatment 10dpi.

As a result of the analysis of ontology and enrichment of differentially expressed genes in the 2dpi treatment, performed in the Agrigo V2 software, in Supplemental Table 1 and in Figure 5, functional categories can be observed, in terms of gene ontology, to which belong the DEG genes that presented annotation in that treatment.

### Specific genes expressed differentially as response of *Vanilla planifolia* Jacks., to *Fusarium*

#### Ribosomal proteins

We found 72 transcripts corresponding to ribosomal proteins that showed a significant increase in their expression patterns in comparison to the control, in fact, the increase in expression arises only at 2dpi, as an early response (Figure 2, Figure 3, figure 5, Figure 4 and Figure 6). Therefore, in 10 dpi did not show significant changes in the expression of Vanilla ribosomal proteins (Figure supplementary. S3 and Figure supplementary. S4). These differentially expressed transcripts correspond to the ribosomal proteins: Ribosomal Protein S5, Ribosomal Protein 5A, Ribosomal Protein S13A, Ribosomal Protein L13, Ribosomal Protein L14, Ribosomal Protein L4 / L1, Sacidicribosomal Protein, Ribosomal Protein L2, S18 Ribosomal Protein, Ribosomal Protein L6, Ribosomal Protein L23AB, Ribosomal Protein Large subunit 16A, Ribosomal ProteinL13, Ribosomal Protein L35Ae, Ribosomal Protein S8, Ribosomal Protein L10, Ribosomal Protein S26e, Ribosomal Protein L5B, Ribosomal Protein SA, Ribosomal Protein S10p / S20e, Ribosomal Protein L18ae / Lx, Ribosomal ProteinL22p / L17, Zinc-Binding Ribosomal Protein, Ribosomal Protein L19e, Glutathione S-transferasem (C-terminal-like, Translation Elongation Factor EF1B / ribosomal protein S6), Ribosomal L29e, Ribosomal S17, Ribosomal Protein L24e, Ribosomal Protein S24e, Ribosomal Protein L36e, Ribosomal Protein L32e, Ribosomal Protein S4, Ribosomal Protein S3Ae, Ribosomal Protein S5 / Elongation Factor G / III / V, Ribosomal Protein L34e, Ribosomal Protein L23 / L15, Ribosomal Protein L30 / L7, Ribosomal Protein L18, Ribosomal Protein L7Ae / L30e / S12e / Gadd45, Ribosomal Protein L18e / L15, Ribosomal Protein S7e, Ribosomal Protein L1p / L10e, Ribosomal Protein S4 (RPS4A), Ribosomal Protein L14p / L23e, Ribosomal Protein S12 / S23, Ribosomal Protein S19e, Ribosomal Protein STV1 (RPL24B, RPL24) Ribosomal Protein L24, Ribosomal Protein S25, Ribosomal Protein S8e, 60S Acidic Ribosomal Protein, Ribosomal Protein S5 Domain 2 Like, Ribosomal L29, Ribosomal Protein L11, Ribosomal L27, Ribosomal L22e. Also, we emphasize that the transcript corresponding to the ribosomal protein S26e is repressed at 2 dpi, while the transcript corresponding to L22e is repressed at 10dpi (Figure 6, Supplemental Table 2).

**Figure 3.**
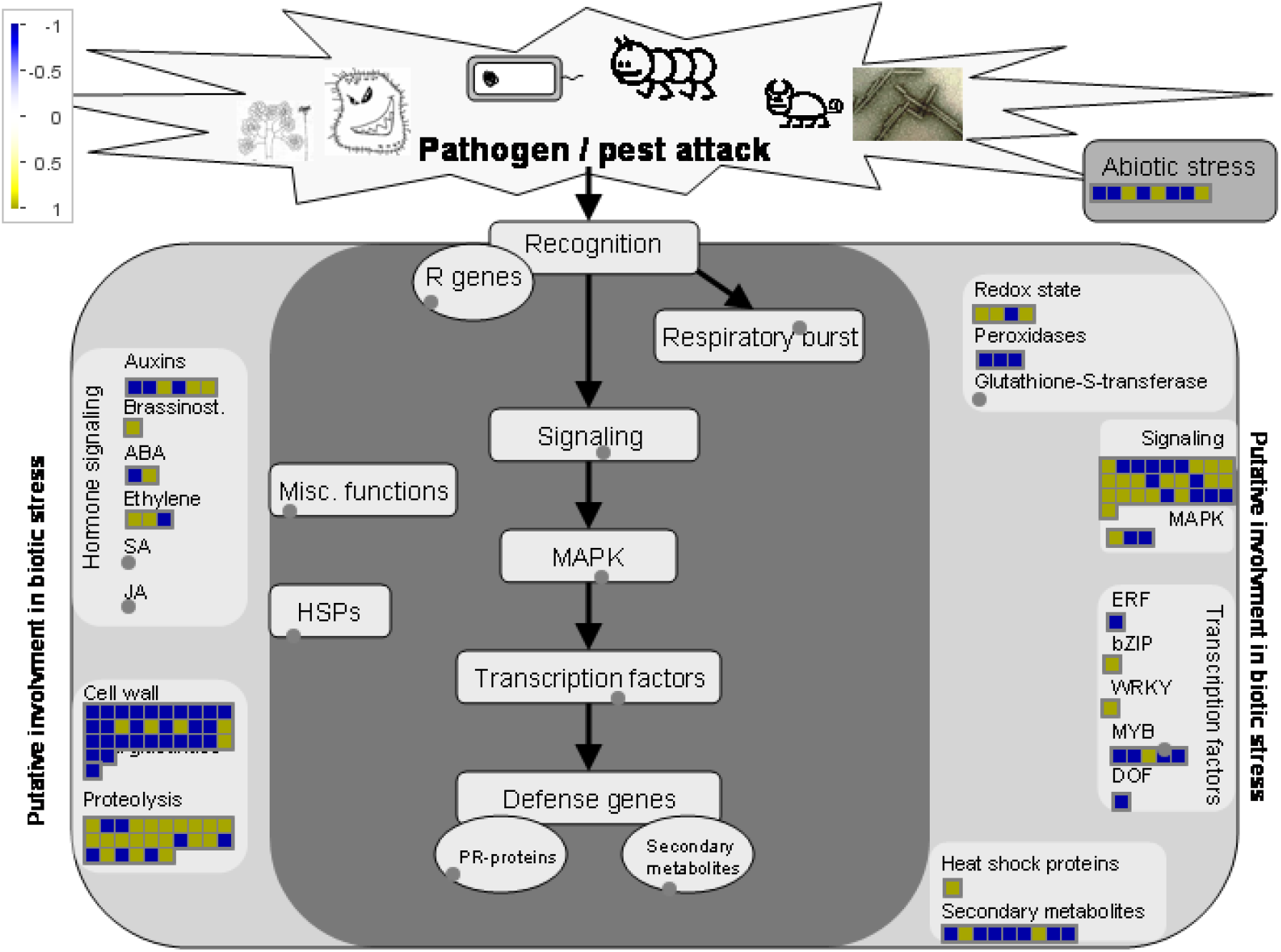
Expression profiles related to biotic stress, in the early response (2dpi) of vanilla to *Fusarium oxysporum* f. sp. *vanillae*. In the present figure, generated with the Mapman software, the expression profiles of the annotated unigenes related to biotic stress in the plants (plant-pathogen interaction), in the early response, to the 2dpi, vanilla ante *Fusarium*.

**Figure 4.**
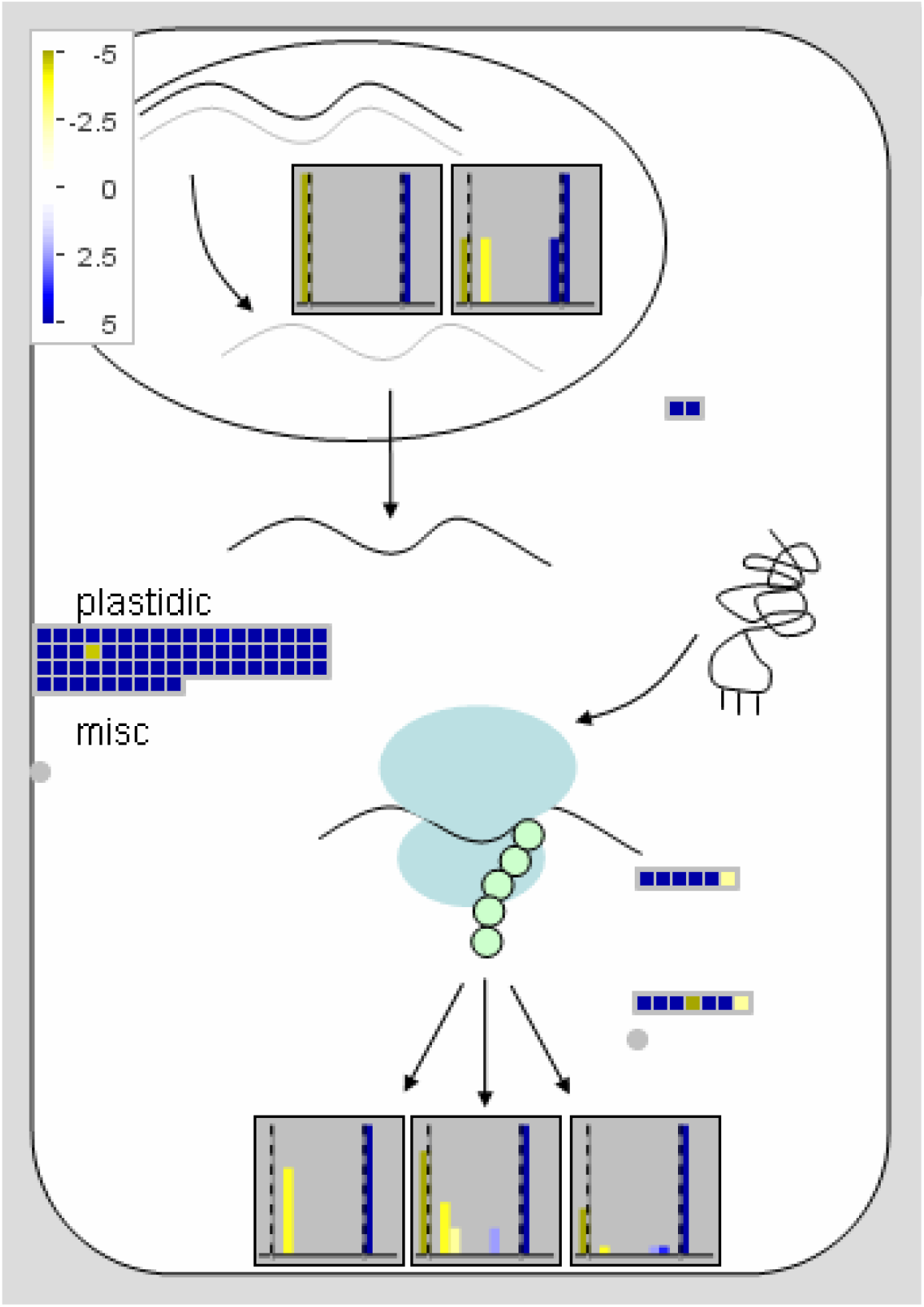
Unigenes involved in the regulation of translation during the early response of vanilla to *Fusarium*. General scheme of translation regulation developed with the Mapman software. The expression profiles are observed, by means of heat maps, of the genes coding for ribosomal proteins, and their participation in the translation.

**Figure 5.**
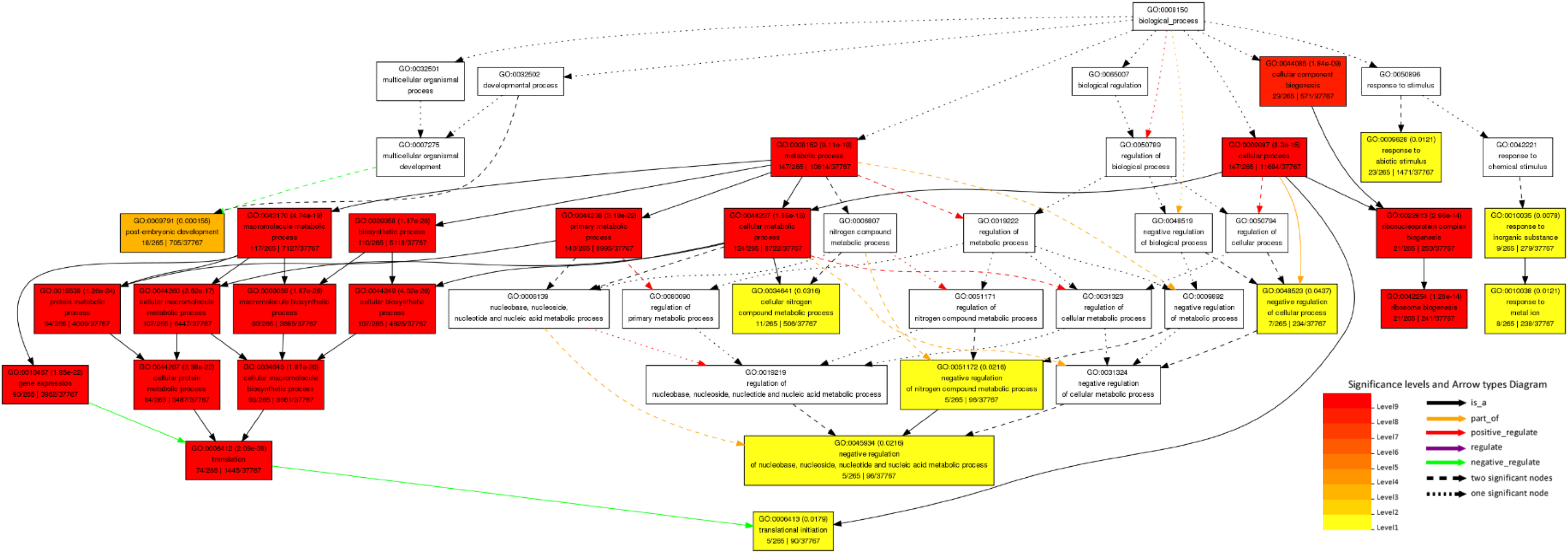
Enrichment of gene ontology terms in the vanilla transcriptome in response to *Fusarium*. Enrichment analysis of gene ontology terms, observing the ontology terms of the Biological Processes category, involved in the vanilla transcriptional response to *Fusarium* at 2dpi.

**Figure 6.**
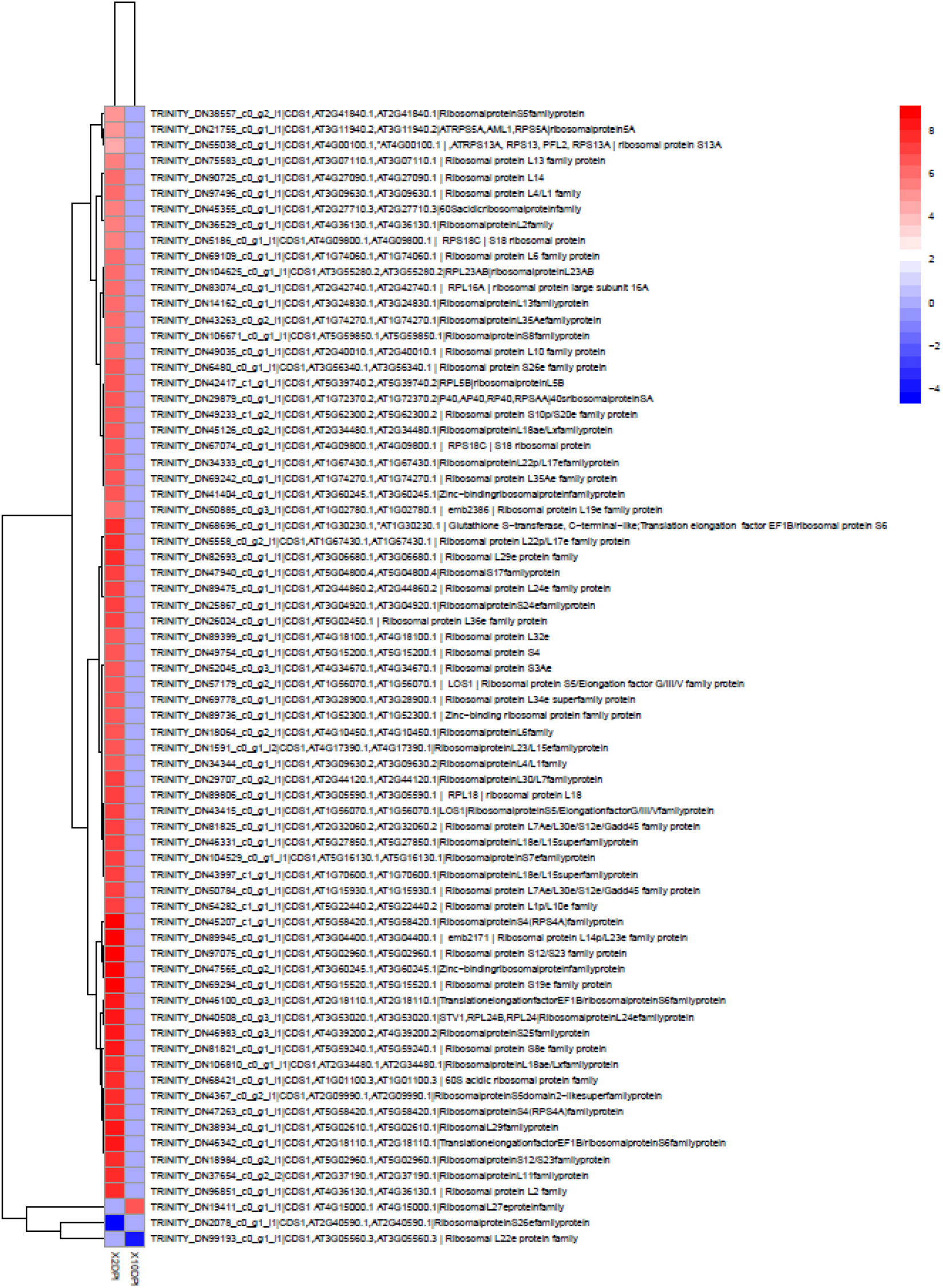
Expression of ribosomal proteins (RPs) as an early vanilla response to *Fusarium* infection. In the present figure we observe the heat map that contrasts the expression of the RPs at 2dpi (right panel) against 10 dpi (left panel), in the vanilla transcriptome in response to *Fusarium*.

**Figure 7.**
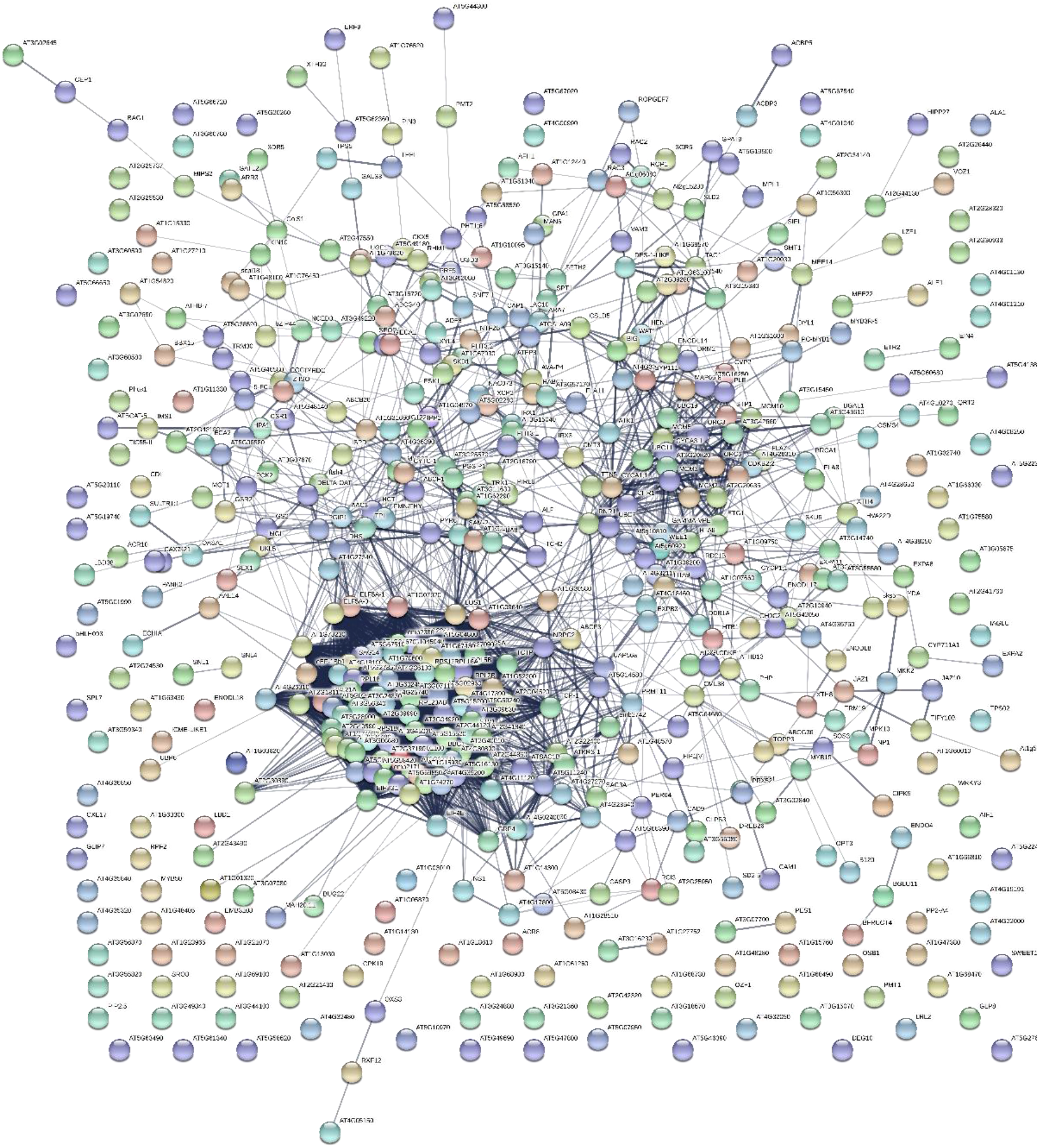
STRING network of genes differentially expressed at 2 dpi in the vanilla response to *Fusarium*. Protein-protein interactions were plotted in STRING by entering the TAIR codes of all genes differentially expressed in the vanilla transcriptome in response to *Fusarium* at 2dpi. The colored bubbles correspond to different proteins present in Arabidopsis. The thickness of the edge corresponds positively to the confidence of the interaction

### Post-embryonic development genes and the transcriptional response of vanilla to *Fusarium*

At 2 dpi we found the expression of a group of 18 genes related to the development of the Arabidopsis embryo, these are listed below: YDA (YODA); SMT1 (STEROL METHYLTRANSFERASE 1); emb2386 (embryo defective 2386); IPP2 (ISOPENTENYL PYROPHOSPHATE: DIMETHYLALLYL PYROPHOSPHATE ISOMERASE 2); MEE22 (MATERNAL EFFECT EMBRYO ARREST 22); GP ALPHA 1 (G PROTEIN ALPHA SUBUNIT 1); emb2171 (embryo defective 2171); BIG (BIG), binding / ubiquitin-protein ligase / zinc ion binding Calossin-like protein required for polar auxin transport BIG (BIG); HPA1 (HISTIDINOL PHOSPHATE AMINOTRANSFERASE 1); NS1, asparagine-tRNA ligase Asparaginyl-tRNA synthetase protein; TAG1 (TRIACYLGLYCEROL BIOSYNTHESIS DEFECT 1); LZF1 (LIGHT-REGULATED ZINC FINGER PROTEIN 1); STV1 (SHORT VALVE1); MEE14 (maternal effect embryo arrest 14) maternal effect embryo arrest 14 (MEE14); SPT1 (SERINE PALMITOYLTRANSFERASE 1); TCTP (TRANSLATIONALLY CONTROLLED TUMOR PROTEIN); TCH2 (TOUCH 2). These genes showed their induction solely at 2pi.

## DISCUSSION

### The early response of Vanilla to *Fusarium*, the RPs and the translational regulation in response to stress

In the present work we found 72 transcripts corresponding to ribosomal proteins, which showed a significant increase in their expression patterns, in response to *Fusarium* infection in vanilla (Figure 6). This increase in expression is only raised at 2dpi, as an early response of the plant to the pathogen. Probably, given the number of transcripts related to ribosomal proteins that show their induction before the invasion of the fungus; the present finding wasn’t documented in a global way, until now. This corresponds to an event of translational regulation as a response to biotic stress, mediated by ribosomal proteins. As reported by Wang et al., in 2013, those who found the global induction of RPs, in Arabidopsis in response to the iron and phosphorus deficit, concluding that, the changes in the expression in the RPs are related to alterations in the composition of the ribosomes and therefore to the translational regulation. In this context, we propose that in the vanilla response to *Fusarium* infection, in the early stages, the expression of the RPs, and therefore the alteration of the ribosome composition, and the translational regulation are key events to understand the response of the plant to the pathogen. As well as to understand the role of PRs and the translational regulation in response to biotic stress.

Now, we emphasize here, the global participation of ribosomal proteins in the response of plants to stress caused by phytopathogenic fungi is evident for the first time; This is consistent with what was reported by Yang et al, in 2013, who observed that the expression of RPL13 is related to the tolerance of potatoes to *Verticullum* and to the induction of the auxin signaling pathway. In the early vanilla response to *Fusarium*, not only did we find the expression of RPL13 (RPS13A, RPL13, RPL34e); but also the expression of RPL10 (Falcone Ferreyra et al, 2010), RPS12 / S23, PRPL19e (Nagaraj et al., 2016), RPs associated with the plant response to bacteria; as well as the expression of RPS6, RPL19, RPL13, RPL7, and RPS2, associated with the plant response to viruses (Yang et al., 2009). Besides, we also find the expression of RPS10, RPS10p / S20e, as reported by Zhang et al, in 2013, in the soy response to *Phytophthora sojae*. The foregoing can be explained as evidence of a process of translational reprogramming in vanilla as specific response of the plant-pathogen interaction.

Moreover, we reported for the first time, the induction of protein expression of: RPS5, RP5A, RPL14, RPL4 / L1, Sacidicribosomal Protein, RPL2, RPS18, RPL6, RPL23aB, RPL16A, RPL35Ae, RPS8, RPS26e, RPL5B, PSA, RPL18ae / Lx, RPL22p / L17, Zinc-Binding Ribosomal PRPL19e, Glutathione S-transferasem (C-terminal-like, EF1B / RBS6), RL29e, RS17, RPL24e, RPS24e, RPL36e, RPL32e, RPS4, RPS3Ae, RPS5 / Elongation Factor G / III / V, RPL34e, RPL23 / L15, RPL30 / L7, RPL18, RPL7Ae / L30e / S12e / Gadd45, RPL18e / L15, RPS7e, RPL1p / L10e, RPS4 (RPS4A), RPL14p / L23e, STV1 (RPL24B, RPL24) RPL24, RPS25, RPS8e, 60S Acidic RP, RPS5 Domain 2 Like, RPL29, RPL11, RPL27, RL22e, as an early plant response to the interaction with phytopathogenic fungi. We highlight here the induction of STV1 (RPL24B, RPL24) RPL24, in the early response of vanilla to *Fusarium*, due to the central role of STV1 in plant development and by its possible participation in the plant response to stress caused by phytopathogenic fungi. Ribosomal proteins, besides constituting the subunits of the ribosome, are essential for their formation and for the decoding of mRNA, as well as for peptides chains to have place. According to Merchante review in 2017, 80 RPs have been identified in Arabidopsis, and notably each of these is encoded, in the Arabidopsis genome, by at least 7 paralogical genes; therefore, these authors and others consider that at least 230 RPs have been identified in the Arabidopsis genome (Barakat et al., 2001). This diversity of genes encoding ribosomal proteins contrasts with the fact that, in general, the presence of a single member of each family of RPs has been observed as part of the subunits of the ribosomes (Perry, 2007).

Given the importance of RPs in the assembly of ribosomes and the translational complex, as well as their central role in the process of mRNA translation, ribosomal proteins are involved in a complex mechanism of translational regulation, as a regulatory mechanism of gene expression at the post-transcriptional level, which has not been elucidated at all, but which recently has received special interest due to its implications in the eukaryotic development and their role in biotic and abiotic stress (Gonskikh and Polacek, 2017; Warner and McIntosh, 2009). In fact, the mutants of Arabidopsis in RPs (RPS18A, RPL24B, RPS5B, RPS13B and RPL27A), “pointed first leaf” mutants, show defective phenotypes related to development, such as the reduction of the growth of shoots and roots, the reduction of the cell proliferation, sensitivity to genotoxins, increase of ploidy in leaf cells, and the presence of denticulate leaves (Revenkova et al., 1999; Ito et al., 2000; Szakonyi and Byrne, 2011; Horiguchi et al., 2012).

In the early vanilla response to *Fusarium* infection, we found the induction of RPS18, RPL24B, RPS5 and RPL27A; so, this could mean the preparation of the vanilla cell for the subsequent translation of genes related to programs of development, growth and cellular proliferation at the root, such as the genes of auxin-mediated signaling and others related to the expression of STV1 (short valve1, RPL24b). This suggest, the presence of a mechanism of translational regulation mediated by vanilla uORFs, and its function in the Vanilla-*Fusarium* interaction, given the relationship of the RPs, and alterations in the composition of the ribosome, with the selective translation of mRNAs containing uORFs, during the development and response to stress (Merchante et al., 2017).

Regarding the expression of RPL10, RPS13 and RPl23aB, and its possible role in the response of vanilla to *Fusarium*, in addition to the fact that the induction of RPL10, RPS13 has already been reported in response to pathogens in plants, it has also been documented that mutation of the Arabidopsis RPL10 gene causes lethality of the female gametophyte; while its overexpression complements this phenotype with the recovery of dwarfism in the mutant of the ACL5 gene, also in Arabidopsis (Imai et al., 2008). In this way, in Arabidopsis RPL10 and RPL10C, they were induced under UV-B stress conditions, a fact that was also observed in maize plants (Casati and Virginia, 2003, Ferreyra et al., 2013). Mutations in RPS13A result in a reduction of cell division, in the retardation of flowering, and in the retardation of the growth of shoots and leaves (Ito et al., 2000).

Other similar phenotypes of growth retardation and fertility reduction have been reported in the RPL23aA mutants, in experiments where the synthesis of the RPL23aA protein is reduced; whereas in the mutant of the RPL23aB gene there are no developmental effects (Degenhardt and Bonham-Smith, 2008). This further evidence, the process of translational reprogramming experienced by the vanilla cell, as a preparation for the subsequent translation of related genes with auxin signaling, development programs and cell growth and proliferation.

The RPs can be regulated by environmental stimuli, like stress or by the signaling of phytohormones. Proof of this is that the transcript levels of RPS15a (and its variants A, C, D and F), in Arabidopsis, increase significantly in response to phytohormones and heat stress (Hulm et al., 2005). This was also evidenced under the treatment with BAP, where an increase in the transcription of RPS14, RPL13, and RPL30 in Arabidopsis was observed (Cherepneva et al., 2003). In the vanilla transcriptome against *Fusarium*, we also find the expression, in addition to the mentioned RPs, the RPL13, RPL14 and RPL30, which indicates their participation, as proteins related to biotic stress and the response mediated by phytohormones, in the vanilla response to the pathogen.

The response to environmental stimuli and ribosomal proteins were also reported in low temperatures conditions, inducing the expression of the RPS6, RPS13 and RPL37 genes in soybean plants (Kim et al, 2004); which has been reported for the homologs RPL13 and C34 in *Brassica* and *E. coli* (Sáez-Vázquez et al., 2000; Tanaka et al., 2001); This fact was also evident in the vanilla transcriptome against *Fusarium*, which indicates the central role of RPL13 in the response to stress in plants, both biotic and abiotic. And as mentioned above, the remarkably overexpression of RPL13 resulted in the tolerance of transgenic potato plants, against the pathogenic fungus *Verticillum dahliae*, with an increase in the expression of defense genes and antioxidant enzymes (Yang et al., 2013).

Here, in the present transcriptome we also found the expression of RPS6, which together with its role in the response to abiotic stress, is also a regulator of translation; therefore, its induction in the present study supports the idea of the translational reprogramming in response to the pathogen. Merchante et al., 2017 described the phosphorylation of RPs, as one of the molecular mechanisms that are part of the complex system of the translational regulation, in which the RPs are involved. So, phosphorylation is involved in plant development and plant stress response (Boex-Fontvieille et al., 2013). The phosphorylation of RPS6 through the TOR signaling pathway is involved in the selective translation of mRNAs (Muench et al., 2012; Merchante et al., 2017). TOR is a highly conserved master coordinator of nutrients, energy and stress signaling in the cell (Xiong and Sheen, 2014). Thus, implying a role of RPs in the direct response to stress, through a specific network of genes related to stress in plants, such as defense proteins, ROS response genes, genes of calcium signaling and the vast network of genes associated with phytohormones (Moin et al., 2016). Finally, given the induction of RPS6 in the vanilla transcriptome in response to *Fusarium*, this may suggest the participation of TOR master regulator in the translational regulation in the Vanilla-*Fusarium* pathosystem.

This represents important clues of the role of the RPs and the variation in their levels of mRNA or protein, as well as the variation of their charge in the polysomes, in the translational regulation. However, if we look at some development events, such as seed germination during the first 24 hours; We found that, there is a drastic increase in transcription and protein synthesis. While, notably the mRNA levels of the RPs remain constant (Merchante et al., 2017). This may mean, there exist a relation between the increase in the synthesis of new RPs, the presence of preexisting mRNAs of RPs, and changes in the efficiency of their translation (Jiménez-López et al., 2011, Merchante et al., 2017). Therefore, in some events such as the cellular response to biotic and abiotic stress, the translation can be decreased, whereas the transcription of alternative RPs increases. So, there is a change in the composition of the ribosome in response to stress.

In addition, that the PRs, like the S6, participates directly in the translational regulation (Morimoto et al., 2002). As we have been describing, it has been widely documented that the RPs have extra-ribosomal functions related to the response to abiotic stress such as salinity, cold and drought in *O. sativa*, where the RPs have been reported: RPS4, RPS7, RPS8, RPS9, RPS10, RPS19, RPS26, RPL2, RPL5, RPL18, RPL44; RPS13, RPS6 and RPL3; while in *G. max* and *B. distachyon* it was reported to RPL27, respectively (Kawasaki et al., 2001, Kim et al., 2004, Bian et al., 2017). In the vanilla transcriptome in response to *Fusarium*, we found most of these RPs, related to abiotic stress, described above, except for RPL2, RPS18 and RPL44, all showing their induction. Given that these proteins are related to the response to abiotic stress and the translational regulation, in the present study we propose a role for these RPs in the general response of plants to stress, or specifically, a participation of these RPs in the response of plants to biotic stress.

Finally, there is growing evidence that RPs not only participate in the translational regulation, but in response to abiotic stress, or regulating translation during development. But they also participate in the response of plants to biotic stress and in particular in the plant response to pathogens, as reported in *N. benthamiana* and *A. thaliana*, where RPL12 and RPL19 participate in the resistance against *P syringae* As well as in *O. sativa* where the differential expression of 27 RPs was reported in response to infection by *Xanthomonas oryzae* (Nagaraj et al., 2016; Saha et al., 2017). In the vanilla transcriptome in response to *Fusarium*, we also observed the induction of RPL12. This is in addition to the other RPs that showed their induction at 2dpi, it also participates in plant development and translational regulation of the response to biotic stress; It is evident that these proteins are induced as an early response of the plant. Therefore, we report their participation in the plant response to biotic stress. Besides, we propose the role of the translational regulation in the early plant response, this in this case of the vanilla, before the attack of pathogens.

### Translational reprogramming in vanilla transcriptome in response to *Fusarium* infection

Stress in plants can even cause a global drop in the translation of proteins, since this process is energetically demanding. However, under stress conditions, a translational regulation mechanism leading to the selection of certain transcripts may also occur. This regulation, mediated by specific genes, may be the key to the survival of plants, as opposed to certain stress conditions, dependent on the expression of a specific group of transcripts (Merchante et al., 2017). Different mechanisms that lead to the global translational regulation have been documented, for example, different stimuli trigger the phosphorylation of RPs and eiFs (Merchante et al., 2017); which results in a contrasting effect in the global translational regulation, which can be repressed or enhanced (up-regulation) (Muench et al., et al 2012; Browning and Bailey-Serres 2015). Regarding the effects of the phosphorylation of the factors of elongation of the translation in the plants, it has been documented that GCN2 (GENERAL CONTROL NONDEREPRESSIBLE2), phosphorylates eIF2α; reducing global protein synthesis, which has implications for growth and development (Zhang et al., 2008, Liu et al., 2015). Notably, in the vanilla transcriptome in response to *Fusarium*, GCN3 was found to be one of the most expressed transcripts, with a logFC of 12. Suggesting the possibly in the early vanilla response to the pathogen, there is a decrease in the translation and the reprogramming of this event, towards the translation of a group of transcripts related to the response of plants to stress. In fact, in the present transcriptome we also observed the induction of eIF2α, the target of GCN2 in the negative regulation of translation, eIF2α has been associated with the response to cold stress in Arabidopsis, since when silenced causes a decrease in the development to the tolerance to stress in the plant. What evidences his participation in the response to stress in plants. In Arabidopsis GCN2 interacts with GCN1 to phosphorylate eIF2α and regulate it negatively; in the vanilla transcriptome in response to *Fusarium*, we found an increase in the expression of GCN1, which associated with GCN3 expression, both translational regulators (Hannig et al., 1990), and the increase in the expression of eIF2α, evidence indirectly the translational reprogramming of proteins, such as an early vanilla response to pathogen attack. Also, in Arabidopsis, the repressor function of GCN1 has been reported, by increasing the formation of polysomes in the *gcn1* mutant, together with the demonstrated role of this gene in the response of Arabidopsis to cold stress (Wang et al, 2017). In addition to the above, the phosphorylation of eIF2α has been related to the negative regulation of translation against purine deficit, UV radiation, cold response, mechanical damage, the response to cadmium and the response to jasmonate, ethylene and salicylic acid (Zhang et al., 208; Sormani et al., 2011; Wang et al, 2017).

### Post-embryonic development genes and the vanilla transcriptional response to *Fusarium*

Sopeña-Torres et al., (2018) reported that YODA, a MAPK3 involved in the establishment of stomata, the development of the embryo, the architecture of the inflorescences and in the development and shape of the lateral organs (Bergmann et al. al., 2004); modulates the immune response of Arabidopsis. By demonstrating how the resistance of the plant to pathogens is compromised, in the *yda* mutants; while the plants that express YDA, show resistance to a broad spectrum of pathogens, such as fungi, bacteria and omicetes with different lifestyles. YDA is part of the same pathway as the Type Kinase Erecta receptor, which together regulate immunity, as a pathway parallel to that regulated by phytohormones and PRRs (Sopeña-Torres et al., 2018). In the present study we observed an increase in the expression of YODA as an early response, at 2dpi, of vanilla before the *Fusarium* infection, as well as an increase in the expression of the SMT1 genes (STEROL METHYLTRANSFERASE 1); emb2386 (embryo defective 2386); IPP2 (ISOPENTENYL PYROPHOSPHATE: DIMETHYLALLYL PYROPHOSPHATE ISOMERASE 2); MEE22 (MATERNAL EFFECT EMBRYO ARREST 22); GP ALPHA 1 (G PROTEIN ALPHA SUBUNIT 1); emb2171 (embryo defective 2171); BIG (BIG), binding / ubiquitin-protein ligase / zinc ion binding Calossin-like protein required for polar auxin transport BIG (BIG); HPA1 (HISTIDINOL PHOSPHATE AMINOTRANSFERASE 1); NS1, asparagine-tRNA ligase Asparaginyl-tRNA synthetase protein; TAG1 (TRIACYLGLYCEROL BIOSYNTHESIS DEFECT 1); LZF1 (LIGHT-REGULATED ZINC FINGER PROTEIN 1); STV1 (SHORT VALVE1); MEE14 (maternal effect embryo arrest 14) maternal effect embryo arrest 14 (MEE14); SPT1 (SERINE PALMITOYLTRANSFERASE 1); TCTP (TRANSLATIONALLY CONTROLLED TUMOR PROTEIN); TCH2 (TOUCH 2). Therefore, we show its role in the early response of Vanilla to the stress caused by the *Fusarium* phytopathogenic fungus. This is also evidence that auxin-mediated signaling and the genes involved in plant development programs also participate in the response to biotic stress and specifically in the plant response to phytopathogenic fungi.

## CONCLUSIONS

In summary, our results demonstrate the global expression of ribosomal proteins associated with the response to stress and development programs, in response to the infection caused by a phytopathogenic fungus in vanilla, which implies a role of RPs in the responses to biotic stress and plant-pathogen interaction.

Our findings further suggest that, the expression of global of the RPs in response to *Fusarium* confirms the alteration of the composition of the ribosomes. Thus, proposing the translational reprogramming as one of the main the plant responses to the attack of the pathogen. This fact, supported by the expression GCN1, GCN3 and eIF2α, key genes in the control of translation in plants.

Lastly, the gene expression related to development and the response to abiotic stress in plants, strongly suggest the change in the conformation of the ribosome and reprogramming, since these genes are related to the expression of different RPs.

## MATERIALS AND METHODS

### Plant material

From plants of *V. planifolia* Jacks., growing in the Totonacapan region (Veracruz, Mexico), samples were collected and propagated under greenhouse conditions. Vigorous and pathogen-free plants were used in the present study at the developmental age of 12 weeks. Such plants exhibited leaf morphology characteristic of *V. planifolia* Jacks., namely elliptic-obtuse in shape and smooth stems. Sixty plants were distributed in three biological replicates. Each biological replicate consisted of groups of 5 plants for each treatment, giving a total of 30 plants for non-treated (control) and treated (experiment) groups. Since two different times were evaluated.

### Infectivity assays

The *in vitro* fungal infection of *V. planifolia* plants was carried out with the JAGH3 strain. This strain of *F. oxysporum* f. sp. *vanillae* has been previously reported as appropriate for bioassays (Adame-García et al., 2012). Briefly, cuttings of *V. planifolia* Jacks., were subjected to darkness during ten days. The absence of light exposition allowed the generation of new roots. A mechanical incision was made in each root under aseptic conditions. Then, roots were exposed to an aqueous solution of spores with a concentration of 1 × 10^6^ CFU of *F. oxysporum* f. sp. *vanillae* (JAGH33 strain). The inoculation was carried out directly on the substrate where cuttings were established. Cuttings belonging to the control group were treated similarly, exposing them to an aqueous solution without spores. Both control and treatment experiments consisted of 30 plants of the same age, established on substrate and maintained under greenhouse conditions with a 12-hour photoperiod (shaded). Sample collection of plant material was carried out in two different periods: 2 and 10 days’ post inoculation (dpi). For each of the treatments and their respective controls, five tissue samples were collected in each case, pooled and processed immediately for RNA extraction. In total, twelve pools were obtained.

### Obtaining total RNA

For the total RNA extraction from the roots of *Vanilla planifolia* Jacks., a protocol was standardized based on what was described by Valderrama-Chairez et al., 2002; which included the extraction of total RNA from 200 mg of root tissue with the help of Trizol Reagent, a treatment with Phenol: chloroform: Isoamyl Alcohol 25: 24: 1, and the subsequent cleaning of the RNA obtained, using silica columns included in the SV Total RNA Isolation System extraction kit from Promega. The integrity of the obtained RNA was determined by electrophoresis in 2% agarose gel, stained with ethidium bromide (EtBr 0.5 ug ml-1) under denaturing conditions. The quality of the total RNA samples was verified by calculating the absorbance ratios A260nm / 280 nm and A260nm / A230nm, obtained for each sample to a UV spectrophotometer (NanoDrop). Finally, the quality and quantity of the RNA was verified, prior to the generation of the cDNA libraries, with the help of the BioAnalyzer, obtaining RIN values close to 8.

### Generation and sequencing of cDNA libraries

The generation and sequencing of the cDNA libraries was carried out in the University Unit of Massive Sequencing and Bioinformatics of the Institute of Biotechnology of the National Autonomous University of Mexico (UUSMB IBT-UNAM). In total, the construction of 12 cDNA libraries was carried out. Afterwards, the sequencing of the cDNA libraries was performed, using the Nextseq 500 illumina platform, of type pair End, with 76 bp with two tags of 8 bp cu, with performances from 5.4 million readings to 10.7 million readings, obtained in total 204 million 517 thousand 080 sequences.

### Assembling the Novo transcriptome of the roots of *Vanilla planifolia* Jacks

Before doing the process of assembly of the transcriptome, a quality analysis was carried out; The analysis of the quality of the readings was made with the FastQC software, and it is an excellent level of quality with all the bases with averages above 32, without the presence of adapters neither in the network of notes, nor in the other side. The bait was made using smalt software 0.7.6 against the reference genus of *Fusarium oxysporum* (*Fusarium oxysporum* f. sp. *lycopersici* 4287), with the identification number of NCBI ASM114995v2. This with the purpose of filtering and discarding the readings corresponding to the pathogen *Fusarium oxysporum*. Then, we took the sequences that had not been used and placed them in “filter” files, in a fast format, using tools and text, and finally with them the set of Novo transcripts was made, using the Trinidad 2.4 software, using the parameters Software To evaluate the quality, the set, obtaining the values N50, ExN50 and L50, using the functions of the Trinity package. Once the set was generated, the completion of the transcripts with the BUSCO software was evaluated. Subsequently, the annotation of the transcriptions was made with the Trinotate software. The search for the open reading frames in the transcriptions was made with the TransDecoder software. Transcripts and amino acid sequences are aligned against the UniProt database using Blastn and Blastx. Using the HMMER software, the presence of PFAM domains in the protein sequences at the time of the transcripts was tested. The annotation of the transcripts was used using Blast2go and the databases of Gene Ontology (GO), KEGG, COG.

### Quantification of results, analysis of differential expression and gene enrichment

To perform the quantification of the transcripts, a method based on the map of the readings can be used within the assembled transcriptome, as part of the Trinidad pipeline using the Bowtie2 software, and later the RSEM. With the results generated by RSEM, with the abundances for each state, by transcript, a table of counts was obtained. The analysis of the differential expression and the integration of the results, has been published through the IDEAMEX website (Jiménez-Jacinto et al, 2019), using the DESeq methods (Anders and Huber, 2010), DESeq2 (Love et al., 2014), EdgeR (Robinson et al., 2013) and NOISeq (Tarazona et al., 2011). To select the differentially expressed genes, we use a padj <= 0.04, FDR <= 0.04 and prob> = 0.96, for DESeq / DESeq2, EdgeR and NOISeq, and a logFC> = 2 in absolute value.

The gene group enrichment analysis (GSEA) was performed using the GO terms in the agriGO v2.0 software. Finally, for the visualization of the transcriptome graphically, for the analysis of genes related to biotic stress, and for the classification of genes. DEG according to each treatment, the ontology of the genes and the expression patterns, in the Mapman V2 software. The heatmaps were generated in software R using the Pheamap and Ggplot2 libraries.

Finally, through the STRING software (http://string.embl.de) the interactions between the gene networks, of the genes differentially expressed at 2 dpi, were analyzed. In order to identify the main gene networks and their interactions, in the early response of vanilla against *Fusarium*.

## LIST OF ABBREVIATIONS

PAMPs: Pathogen-Associated Molecular Patterns.
RNA-Seq: (high-throughput) sequencing of RNA.
PTI: PAMP-Triggered Immunity;
ETI: Effector-Triggered Immunity
PRRs: Plant Pattern Recognition Receptors
MAPKs: Mitogen-Activated Protein Kinases
HR: Hypersensitive Response
2 dpi: 2 days after inoculation
10 dpi: 10 days after inoculation
DEG: Differentially Expressed Genes
RP: Ribosomal Protein
RPS: Small Subunit Ribosomal Protein
RPL: Large Subunit Ribosomal Protein
YDA: YODA
SMT1: STEROL METHYLTRANSFERASE 1
emb2386: embryo defective 2386
IPP2: ISOPENTENYL PYROPHOSPHATE DIMETHYLALLYL PYROPHOSPHATE ISOMERASE 2
MEE22: MATERNAL EFFECT EMBRYO ARREST 22
GP ALPHA 1: G PROTEIN ALPHA SUBUNIT 1
emb2171: embryo defective 2171
BIG: Binding / ubiquitin-protein ligase / zinc ion binding Calossin-like protein required for polar auxin transport BIG
HPA1: HISTIDINOL PHOSPHATE AMINOTRANSFERASE 1
NS1: asparagine-tRNA ligase Asparaginyl-tRNA synthetase protein
TAG1: TRIACYLGLYCEROL BIOSYNTHESIS DEFECT 1
LZF1: LIGHT-REGULATED ZINC FINGER PROTEIN 1
STV1: SHORT VALVE1
MEE14: maternal effect embryo arrest 14; maternal effect embryo arrest 14 (MEE14)
SPT1: SERINE PALMITOYLTRANSFERASE 1
TCTP: TRANSLATIONALLY CONTROLLED TUMOR PROTEIN
TCH2: TOUCH 2
eiFs: Eukaryotic Initiation Factors
CGN: GENERAL CONTROL NONDEREPRESSIBLE
logFC: log2 of fold change
MAPK: Mitogen-Activated Protein Kinases

## DECLARATIONS

### Ethics approval and consent to participate

Not applicable

### Consent for publication

Not applicable

### Availability of data and material

The datasets used and/or analyzed during the current study are available from the corresponding author on reasonable request.

### Competing interests

The authors declare that they have no competing interests

### Funding

The funds for the realization of the present investigation were provided by Tecnológico Nacional de México, under the program and / or call ... The design of the present project, as well as its execution, the collection of the data and its analysis and interpretation, was the responsibility of the authors. The bioassays, the experimental phase was carried out in the Laboratory of Microbial Interactions, of the Faculty of Agricultural Sciences of the Universidad Veracruzana.

### Author’s contributions

All authors contributed to the research project design and manuscript preparation. Conceived and designed the experiments MTSC, LIA, JAG and MLR. Performed the experiments MTSC, EEEH. Analyzed the data VJJ, LVA, MTSC, EEEH and JGJ. Wrote the paper: MTSC, EEEH, JGJ, VJJ and LVA (with input from the other authors). All authors read and approved the final manuscript.

## Acknowledgements

To the Consejo Nacional de Ciencia y Tecnología (CONACYT), for the scholarship granted to the author to carry out his postgraduate studies. To the Tecnológico Nacional de México for the financing granted to carry out the present investigation. To the Biologist Lolvin Delaurens-Santacruz, to Dr. Alejandro Blanco Labra and to Dr. Gastón Jiménez-Contreras, for their valuable help in the revision of this text. To Dr. Matías Baranzelli for his valuable help in the structuring and revision of the manuscript. To Edder Darío Aguilar-Méndez, Daniel Abisaí Jerez-Prieto, Dulce Natali Gómez-Hernández and Cecilio Mauricio-Ramos, for their technical assistance during RNA extractions.

