## Supplemental Figures and Tables for "Increase in ribosomal proteins activity: Translational reprogramming in *Vanilla planifolia* Jacks., against *Fusarium* infection"

**SUPLEMENTARY MATERIAL**

**1) Comparative analysis of the different differential expression methods applied to the transcriptome of *Vanilla planifolia* Jacks.**

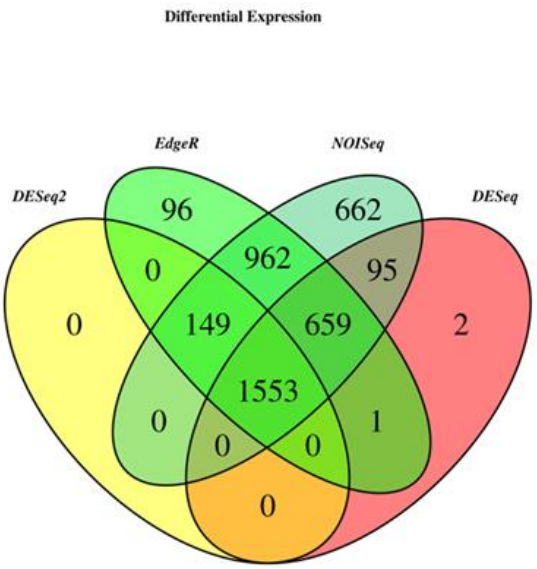


**2dpi**

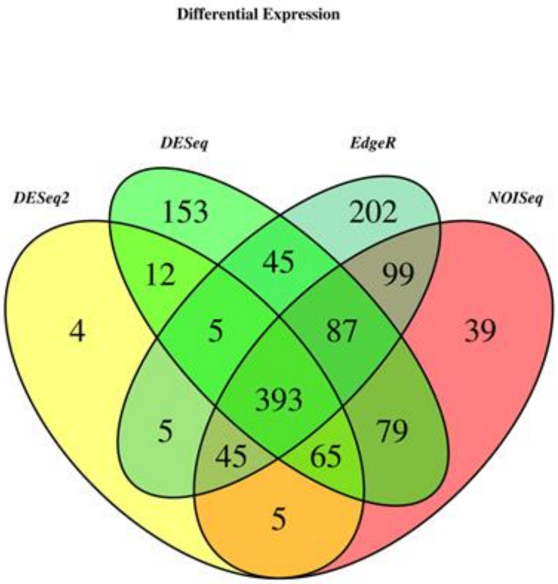


**10dpi**

**Supplemental Figure S1.** Venn diagram showing the comparison of the differentially expressed unigenes obtained with the methods DESeq2, EdgeR, NOISeq, and DESeq. At the center of the diagram we observed that the EdgeR method comprises the great majority of genes determined by the other methods. The right panel corresponds to 2 dpi, while the left panel corresponds to 10 dpi.

**2) Global expression profiles in response to infection caused by *Fusarium oxysporum* f. sp. *vanillae* in vanilla.**

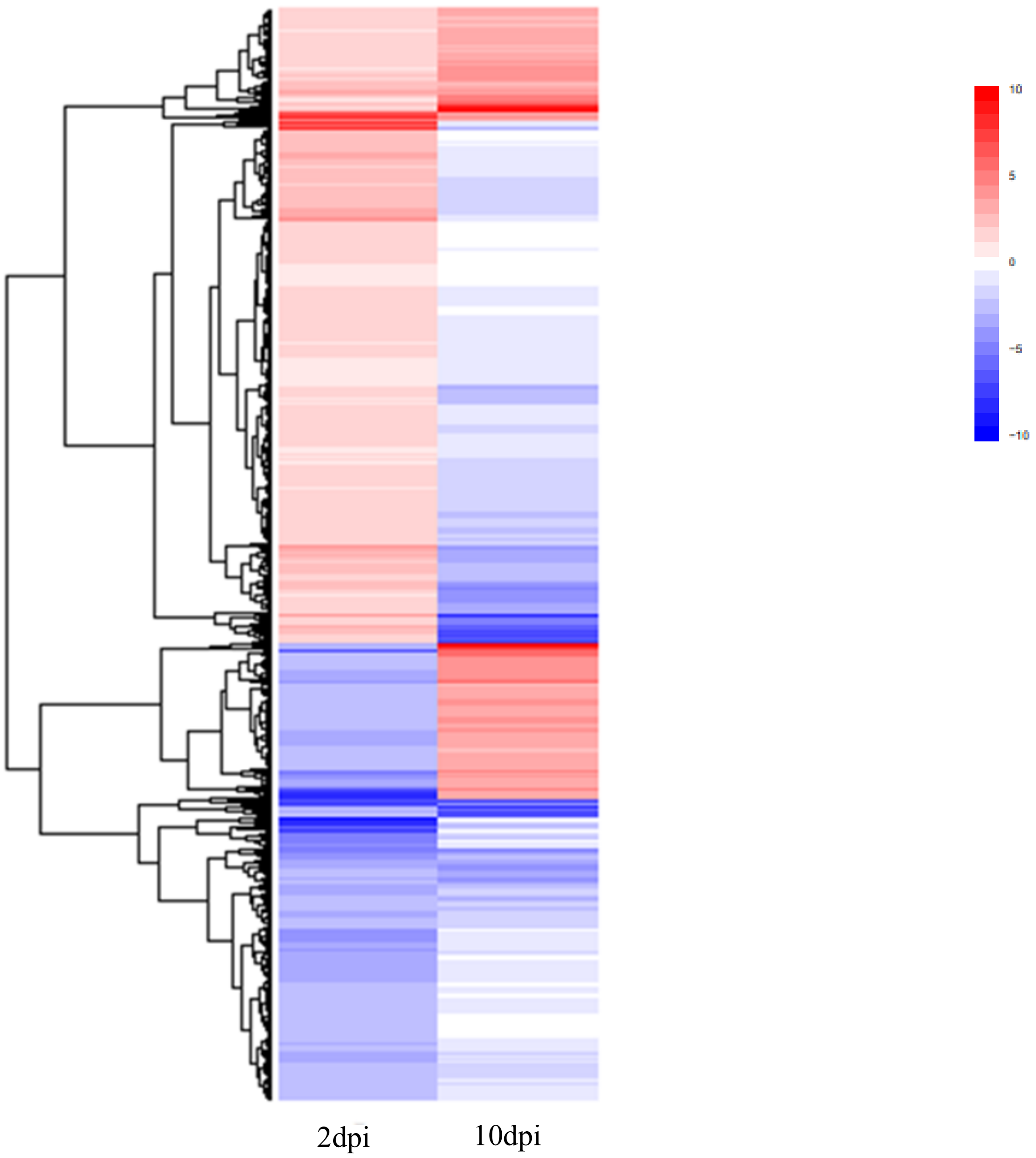


**Supplemental Figure S2.** Heat map that contrasts the global vanilla response to *Fusarium oxysporum* f. sp. *vanillae*. On the right we observe the early response (2dpi); while in the left panel it presents the response to 10dpi. All differentially expressed unigenes are included.

**3) Expression profiles related to biotic stress, in the late response (10dpi) of vanilla to *Fusarium oxysporum* f. sp. *vanillae***

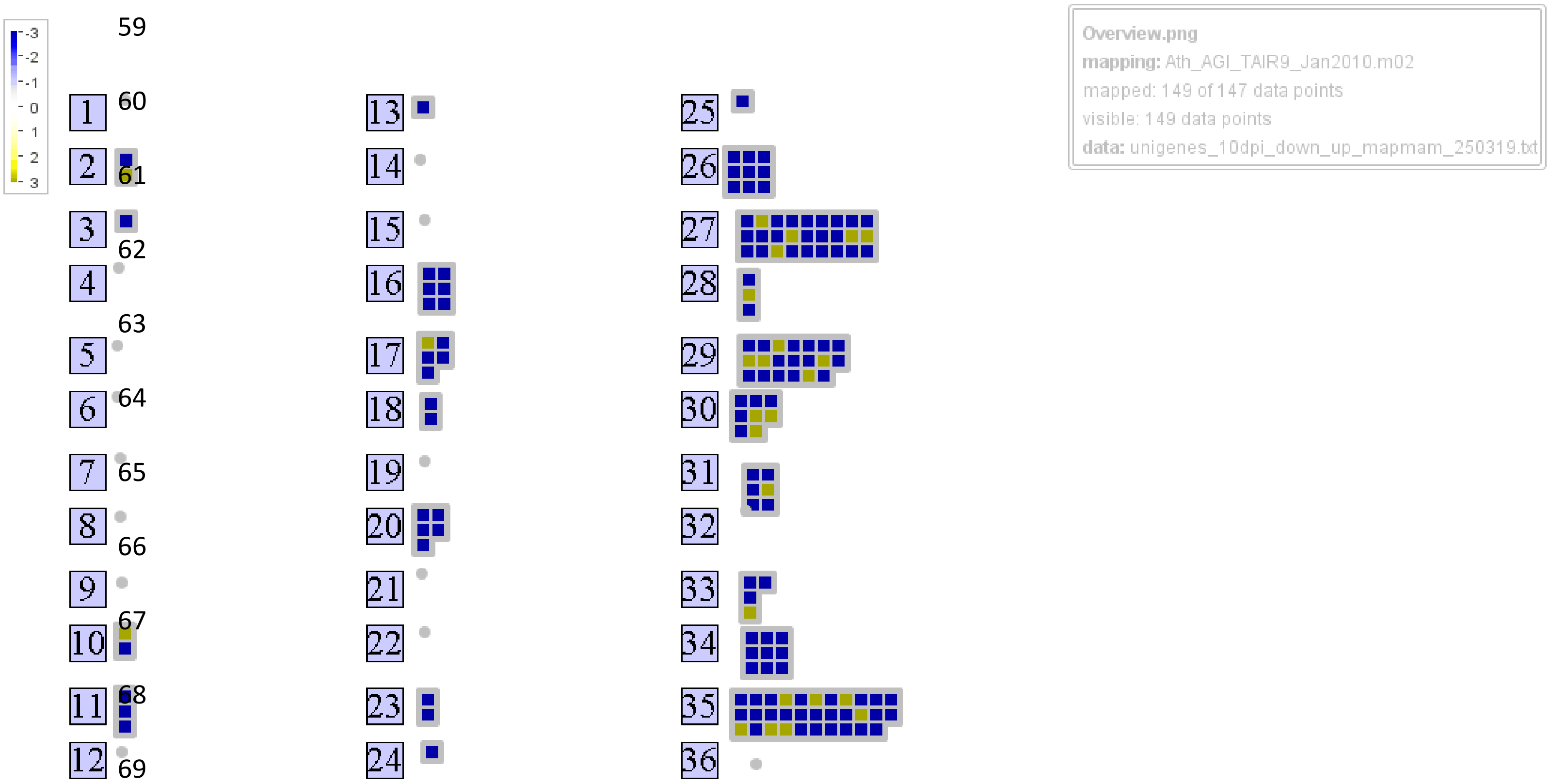


**Supplemental Figure S3.** Heat map indicating the expression profiles of the annotated DEG unigenes, corresponding to 10dpi. The numbers in the figure correspond to different categories of gene ontology, as described below: 25 C1-metabolism, 11 lipid metabolism, 3 minor CHO metabolism, 13 amino acid metabolism, 16 secondary metabolism, 26 misc, 17 hormone metabolism, 30 signalling, 31 cell, 23 nucleotide metabolism, 27 RNA, 28 DNA, 33 development, 24 Biodegradation of Xenobiotics, 18 Co-factor and vitamine metabolism, 35 not assigned, 34 transport, 29 protein, 20 stress, 2 major CHO metabolism, 10 cell wall.

**4) Expression profiles related to biotic stress, in the late response (10dpi) of vanilla to *Fusarium*.**

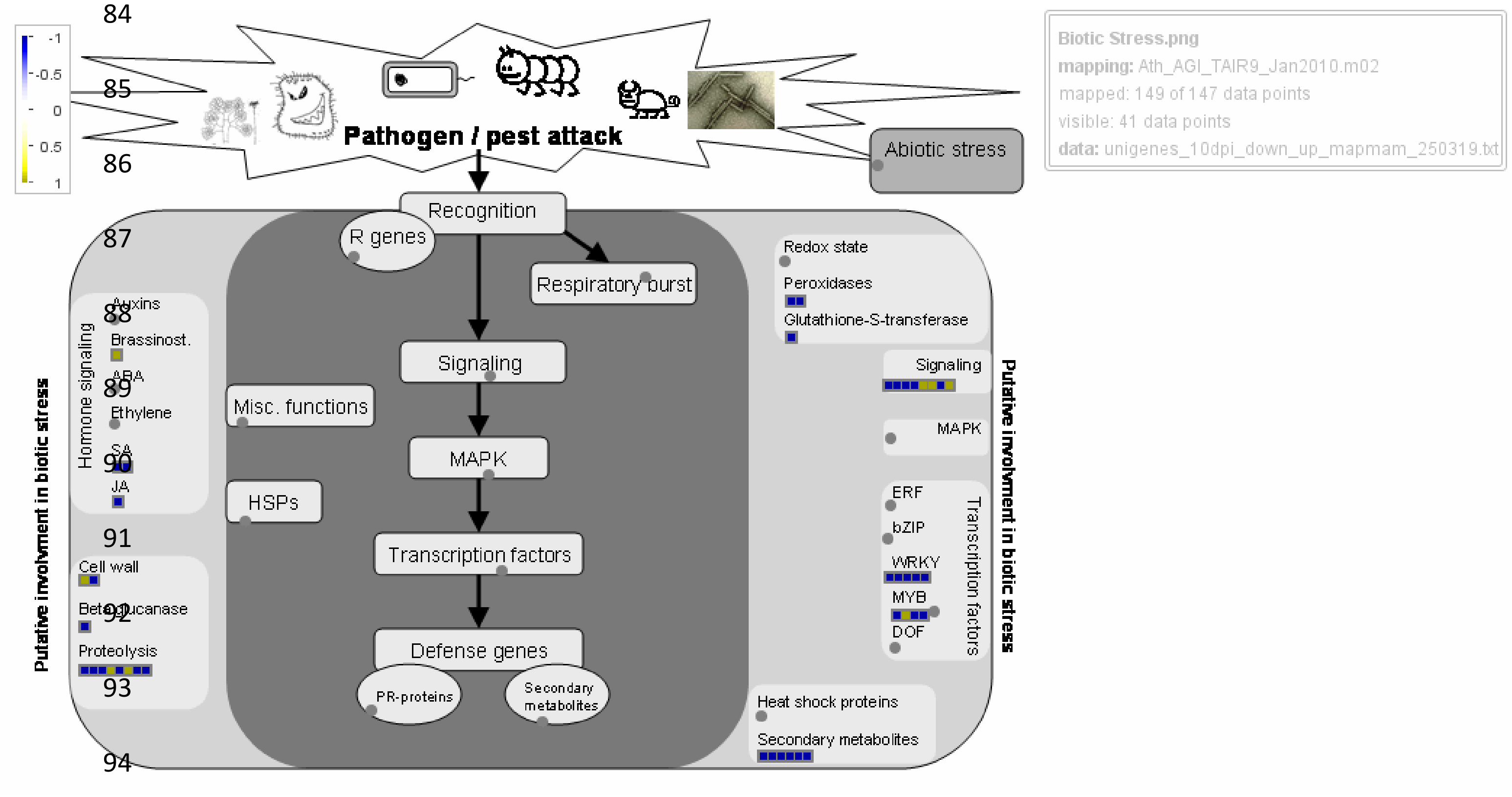


**Supplemental Figure S4.** In the present figure, generated with the Mapman software, the expression profiles of the annotated unigenes related to biotic stress in the plants (plant-pathogen interaction), in the late response, to the 10dpi, of vanilla before *Fusarium*.

5) **STRING network of genes differentially expressed at 10 dpi in the vanilla response to *Fusarium*.**

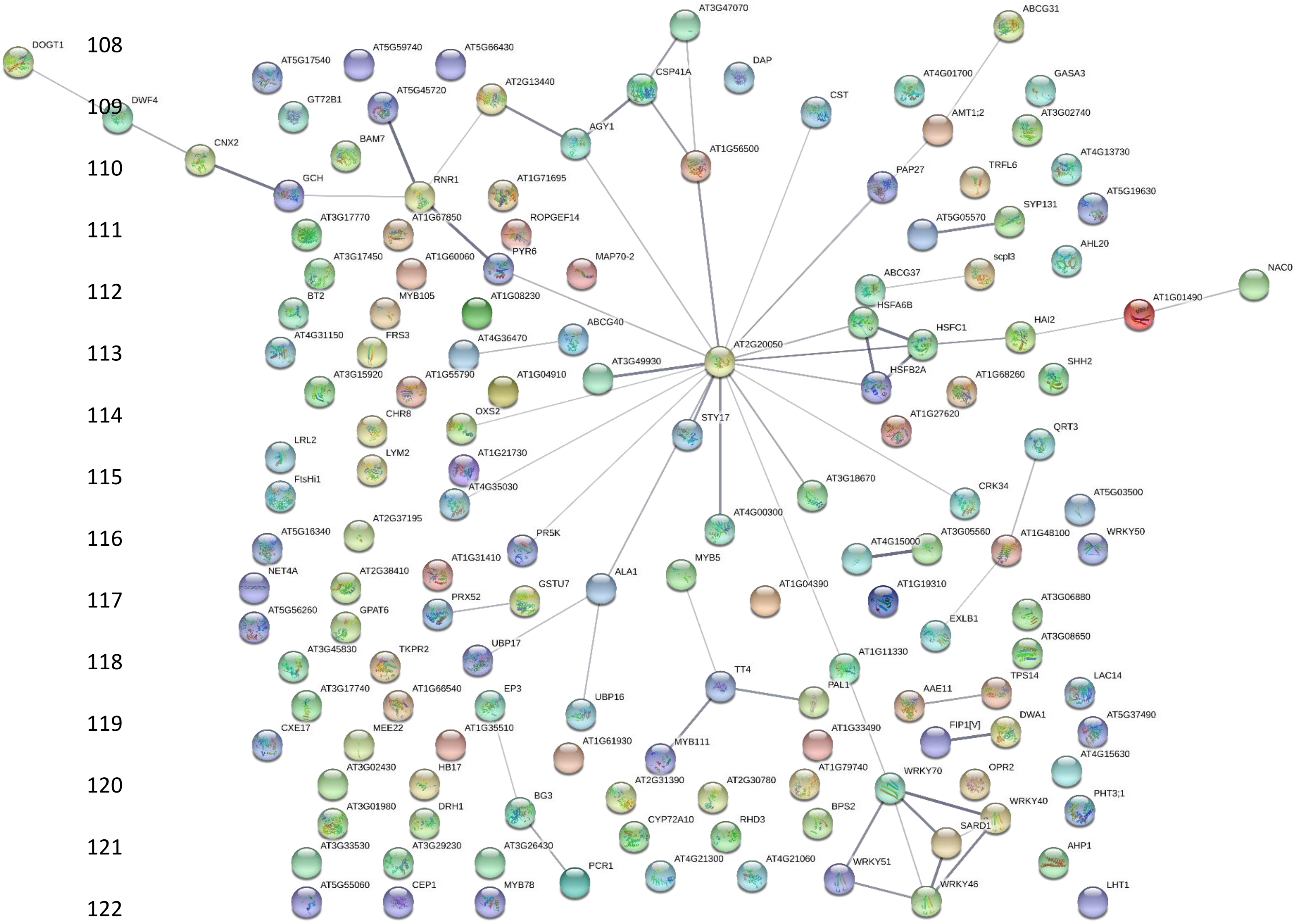


**Supplemental Figure S5.** Protein-protein interactions were plotted in STRING by entering the TAIR codes of all genes differentially expressed in the vanilla transcriptome in response to *Fusarium* at 10 dpi. The colored bubbles correspond to different proteins present in Arabidopsis. The thickness of the edge corresponds positively to the confidence of the interactio

**1) Main categories of gene ontology corresponding to the transcripts of the early response (2 dpi) of vanilla before *Fusarium*.**

| GO term | Ontology | Description | Number | Reference | p-value | FDR |
|---|---|---|---|---|---|---|
| GO:0006412 | P | translation | 74 | 1445 | 2.90E-41 | 2.10E-38 |
| GO:0034645 | P | Celular macromolecule biosynthetic process | 93 | 3661 | 6.50E-29 | 1.90E-26 |
| GO:0009058 | P | biosynthetic process | 110 | 5118 | 7.90E-29 | 1.90E-26 |
| GO:0009059 | P | macromolecule biosynthetic process | 93 | 3685 | 1.00E-28 | 1.90E-26 |
| GO:0044249 | P | cellular biosynthetic process | 107 | 4925 | 2.80E-28 | 4.00E-26 |
| GO:0019538 | P | protein metabolic process | 94 | 4009 | 1.10E-26 | 1.30E-24 |
| GO:0010467 | P | gene expression | 90 | 3962 | 1.80E-24 | 1.90E-22 |
| GO:0044267 | P | cellular protein metabolic process | 84 | 3487 | 2.70E-24 | 2.40E-22 |
| GO:0044238 | P | primary metabolic process | 140 | 8995 | 4.00E-24 | 3.20E-22 |
| GO:0043170 | P | macromolecule metabolic process | 117 | 7127 | 6.60E-21 | 4.70E-19 |
| GO:0008152 | P | metabolic process | 147 | 10614 | 1.40E-20 | 9.10E-19 |
| GO:0044260 | P | cellular macromolecule metabolic process | 107 | 6447 | 4.40E-19 | 2.60E-17 |
| GO:0044237 | P | cellular metabolic process | 124 | 8722 | 3.40E-17 | 1.90E-15 |
| GO:0009987 | P | cellular process | 147 | 11684 | 1.60E-16 | 8.30E-15 |
| GO:0042254 | P | ribosome biogenesis | 21 | 241 | 2.70E-16 | 1.30E-14 |
| GO:0022613 | P | ribonucleoprotein complex biogenesis | 21 | 253 | 6.60E-16 | 3.00E-14 |
| GO:0044085 | P | cellular component biogenesis | 23 | 571 | 4.40E-11 | 1.80E-09 |
| GO:0003735 | F | structural constituent of ribosome | 63 | 494 | 4.20E-57 | 1.40E-54 |

| | | | | | | |
|---|---|---|---|---|---|---|
| GO:0005198 | F | structural molecule activity | 63 | 659 | 5.20E-50 | 8.30E-48 |
| GO:0008135 | F | translation factor activity, nucleic acid binding | 11 | 181 | 1.20E-07 | 1.30E-05 |
| GO:0003746 | F | translation elongation factor activity | 6 | 38 | 6.10E-07 | 4.90E-05 |
| GO:0022626 | C | cytosolic ribosome | 62 | 336 | 4.60E-65 | 1.00E-62 |
| GO:0044445 | C | cytosolic part | 58 | 360 | 1.00E-57 | 1.10E-55 |
| GO:0033279 | C | ribosomal subunit | 58 | 389 | 5.50E-56 | 4.10E-54 |
| GO:0005840 | C | ribosome | 63 | 524 | 1.20E-55 | 6.50E-54 |
| GO:0030529 | C | ribonucleoprotein complex | 63 | 703 | 2.00E-48 | 8.80E-47 |
| GO:0043232 | C | intracellular non-membrane-bounded organelle | 69 | 1040 | 3.70E-45 | 1.20E-43 |
| GO:0043228 | C | non-membrane-bounded organelle | 69 | 1040 | 3.70E-45 | 1.20E-43 |
| GO:0005829 | C | cytosol | 64 | 912 | 3.90E-43 | 1.10E-41 |
| GO:0022625 | C | cytosolic large ribosomal subunit | 36 | 162 | 3.20E-40 | 7.70E-39 |
| GO:0015934 | C | large ribosomal subunit | 36 | 225 | 1.10E-35 | 2.40E-34 |
| GO:0044422 | C | organelle part | 80 | 2562 | 3.30E-30 | 6.00E-29 |
| GO:0044446 | C | intracellular organelle part | 80 | 2561 | 3.20E-30 | 6.00E-29 |
| GO:0032991 | C | macromolecular complex | 70 | 2180 | 7.00E-27 | 1.20E-25 |
| GO:0022627 | C | cytosolic small ribosomal subunit | 22 | 130 | 1.20E-22 | 1.80E-21 |
| GO:0015935 | C | small ribosomal subunit | 22 | 164 | 1.10E-20 | 1.60E-19 |
| GO:0044444 | C | cytoplasmic part | 106 | 6289 | 2.30E-19 | 3.20E-18 |
| GO:0005737 | C | cytoplasm | 109 | 6822 | 2.80E-18 | 3.70E-17 |
| GO:0005730 | C | nucleolus | 20 | 209 | 2.70E-16 | 3.20E-15 |

| | | | | | | |
|---|---|---|---|---|---|---|
| GO:0043229 | C | intracellular organelle | 112 | 8149 | 4.40E-14 | 4.90E-13 |
| GO:0005622 | C | intracellular | 125 | 9671 | 4.40E-14 | 4.90E-13 |
| GO:0043226 | C | organelle | 112 | 8155 | 4.70E-14 | 4.90E-13 |
| GO:0031981 | C | nuclear lumen | 22 | 374 | 1.00E-13 | 1.00E-12 |
| GO:0044424 | C | intracellular part | 120 | 9302 | 2.70E-13 | 2.60E-12 |
| GO:0043233 | C | organelle lumen | 24 | 539 | 2.20E-12 | 1.90E-11 |
| GO:0070013 | C | intracellular organelle lumen | 24 | 539 | 2.20E-12 | 1.90E-11 |
| GO:0044428 | C | nuclear part | 24 | 543 | 2.50E-12 | 2.10E-11 |
| GO:0031974 | C | membrane-enclosed lumen | 24 | 546 | 2.80E-12 | 2.30E-11 |
| GO:0044464 | C | cell part | 152 | 15217 | 1.80E-08 | 1.30E-07 |
| GO:0005623 | C | cell | 152 | 15217 | 1.80E-08 | 1.30E-07 |
| GO:0016020 | C | membrane | 56 | 4068 | 7.70E-07 | 5.60E-06 |

**Supplemental Table 1.** List of the main functional categories, determined with the AgriGO 2.0 software, to which the annotated transcripts belong, which show differential expression at 2 dpi, in the vanilla transcriptome in response to *Fusarium*

2) **Top 100 list of transcripts expressed (up-regulated) at 2 dpi in the vanilla transcriptome, in response to *Fusarium*.**

| **Name** | logFc |
|---|---|
| TRINITY_DN89410_c0_g1_i1\|CDS1,AT1G64550.1,"AT1G64550.1 \| ATGCN3, GCN3 \| general control non-repressible 3 | 12.7703632 |
| TRINITY_DN104321_c0_g1_i1\|CDS1 AT5G36950.1 AT5G36950.1\|DegP10\|DegPprotease10 | 11.1087838 |
| TRINITY_DN83426_c0_g1_i1\|CDS1,AT1G18610.1,AT1G18610.1 \| Galactose oxidase/kelch repeat superfamily protein | 10.4627113 |
| TRINITY_DN54807_c0_g2_i1\|CDS1,AT5G60390.3,AT5G60390.3 \| GTP binding Elongation factor Tu family protein | 10.1980704 |
| TRINITY_DN6978_c0_g1_i1\|CDS1,AT1G69100.1,AT1G69100.1 \| Eukaryotic aspartyl protease family protein | 10.1743963 |
| TRINITY_DN63794_c0_g1_i3\|CDS1, AT3G22590.1,"AT3G22590.1 \| PHP, CDC73 \| PLANT HOMOLOGOUS TO PARAFIBROMIN | 10.014206 |
| TRINITY_DN75904_c0_g1_i1\|CDS1,AT4G00680.1,AT4G00680.1 \| ADF8 \| actin depolymerizing factor 8 | 9.88375811 |
| TRINITY_DN91577_c0_g1_i1\|CDS1,AT5G66720.1,AT5G66720.1 \| Protein phosphatase 2C family protein | 9.82955438 |
| TRINITY_DN29703_c0_g1_i2\|CDS1 AT4G00900.1 AT4G00900.1\|ECA2,ATECA2\|ER-typeCa2+-ATPase2 | 9.64577056 |
| TRINITY_DN66086_c0_g1_i2\|CDS1,AT1G10760.1,"AT1G10760.1 \| SEX1, SOP1, SOP, GWD1, GWD \| Pyruvate phosphate dikinase, PEP/pyruvate binding domain | 9.57132483 |
| TRINITY_DN65143_c0_g1_i1\|CDS1,AT4G01210.1,AT4G01210.1 \| glycosyl transferase family 1 protein | 9.34292421 |
| TRINITY_DN90514_c0_g1_i1\|CDS1,AT3G02260.1,"AT3G02260.1 \| BIG, DOC1, TIR3, UMB1, ASA1, LPR1, CRM1 \| auxin transport protein (BIG) | 9.30763592 |
| TRINITY_DN65916_c0_g1_i2\|CDS1,AT1G10095.1,AT1G10095.1 \| Protein prenylyltransferase superfamily protein | 9.28687646 |
| TRINITY_DN18150_c0_g1_i1\|CDS1 AT3G15720.1 AT3G15720.1\|Pectinlyase-likesuperfamilyprotein | 9.21071873 |
| TRINITY_DN45207_c1_g1_i1\|CDS1 AT5G58420.1 AT5G58420.1\|RibosomalproteinS4(RPS4A)familyprotein | 9.13507978 |
| TRINITY_DN74463_c0_g1_i1\|CDS1,AT4G16370.1,"AT4G16370.1 \| ATOPT3, OPT3 \| oligopeptide transporter | 9.11594149 |
| TRINITY_DN10093_c0_g2_i1\|CDS1 AT1G06080.1 AT1G06080.1\|ADS1\|delta9desaturase1 | 9.10695891 |
| TRINITY_DN89649_c0_g1_i1\|CDS1,AT4G35840.1,AT4G35840.1 \| RING/U-box superfamily protein | 9.07751021 |
| TRINITY_DN97412_c0_g1_i1\|CDS1,AT1G18610.1,AT1G18610.1 \| Galactose oxidase/kelch repeat superfamily protein | 9.02973479 |
| TRINITY_DN64271_c0_g1_i1\|CDS1,AT2G24820.1,AT2G24820.1 \| TIC55-II \| translocon at the inner envelope membrane of chloroplasts 55-II | 9.0048794 |
| TRINITY_DN89945_c0_g1_i1\|CDS1,AT3G04400.1,AT3G04400.1 \| emb2171 \| Ribosomal protein L14p/L23e family protein | 8.93054813 |

| | |
|---|---|
| TRINITY_DN41912_c0_g2_i1\|CDS1 AT1G13950.1 AT1G13950.1\|EIF-5A,ELF5A-1,ATELF5A-1,EIF5A\|eukaryoticelongationfactor5A-1 | 8.91277437 |
| TRINITY_DN97075_c0_g1_i1\|CDS1,AT5G02960.1,AT5G02960.1 \| Ribosomal protein S12/S23 family protein | 8.88831654 |
| TRINITY_DN47565_c0_g2_i1\|CDS1 AT3G60245.1 AT3G60245.1\|Zinc-bindingribosomalproteinfamilyprotein | 8.83080173 |
| TRINITY_DN69294_c0_g1_i1\|CDS1,AT5G15520.1,AT5G15520.1 \| Ribosomal protein S19e family protein | 8.74353089 |
| TRINITY_DN66456_c0_g1_i5\|CDS1,AT2G34780.1,"AT2G34780.1 \| EMB1611, MEE22 \| maternal effect embryo arrest 22 | 8.71800884 |
| TRINITY_DN48558_c0_g1_i2\|CDS1,AT2G22240.2,"AT2G22240.2 \| ATMIPS2, MIPS2, ATIPS2 \| myo-inositol-1-phosphate synthase 2 | 8.7029331 |
| TRINITY_DN21909_c0_g1_i1\|CDS1 AT5G59300.1 AT5G59300.1\|UBC7,ATUBC7\|ubiquitincarrierprotein7 | 8.66463863 |
| TRINITY_DN3969_c0_g1_i1\|CDS1 AT1G61950.1 AT1G61950.1\|CPK19\|calcium-dependentproteinkinase19 | 8.60365732 |
| TRINITY_DN40508_c0_g3_i1\|CDS1 AT3G53020.1 AT3G53020.1\|STV1,RPL24B,RPL24\|RibosomalproteinL24efamilyprotein | 8.56960655 |
| TRINITY_DN46983_c0_g3_i1\|CDS1 AT4G39200.2 AT4G39200.2\|RibosomalproteinS25familyprotein | 8.56637826 |
| TRINITY_DN39408_c0_g1_i1\|CDS1 AT3G49010.3 AT3G49010.3\|ATBBC1,BBC1,RSU2\|breastbasicconserved1 | 8.53411765 |
| TRINITY_DN65733_c0_g1_i1\|CDS1,AT5G08430.1,AT5G08430.1 \| SWIB/MDM2 domain;Plus-3;GYF | 8.52564742 |
| TRINITY_DN105536_c0_g1_i1\|CDS1 AT2G43180.2 AT2G43180.2\|Phosphoenolpyruvatecarboxylasefamilyprotein | 8.50148232 |
| TRINITY_DN66439_c0_g1_i4\|CDS1,AT1G01320.2,AT1G01320.2 \| Tetratricopeptide repeat (TPR)-like superfamily protein | 8.49327629 |
| TRINITY_DN48566_c0_g2_i1\|CDS1,AT4G28390.1,"AT4G28390.1 \| AAC3, ATAAC3 \| ADP/ATP carrier 3 | 8.47172029 |
| TRINITY_DN97500_c0_g1_i1\|CDS1,AT2G21430.1,AT2G21430.1 \| Papain family cysteine protease | 8.44452373 |
| TRINITY_DN63556_c0_g1_i2\|CDS1,AT2G22400.1,AT2G22400.1 \| S-adenosyl-L-methionine-dependent methyltransferases superfamily protein | 8.4025303 |
| TRINITY_DN46100_c0_g3_i1\|CDS1 AT2G18110.1 AT2G18110.1\|TranslationelongationfactorEF1B/ribosomalproteinS6familyprotein | 8.40071786 |
| TRINITY_DN75225_c0_g1_i1\|CDS1,AT5G49180.1,AT5G49180.1 \| Plant invertase/pectin methylesterase inhibitor superfamily | 8.38617923 |
| TRINITY_DN85729_c0_g1_i1\|CDS1,AT4G04930.1,AT4G04930.1 \| DES-1-LIKE \| fatty acid desaturase family protein | 8.35016149 |
| TRINITY_DN23511_c0_g1_i1\|CDS1 AT2G46680.2 AT2G46680.2\|ATHB-7,ATHB7,HB-7\|homeobox7 | 8.33093044 |
| TRINITY_DN18858_c0_g1_i1\|CDS1 AT5G35630.3 AT5G35630.3\|GS2,GLN2,ATGSL1\|glutaminesynthetase2 | 8.25844818 |
| TRINITY_DN18858_c0_g1_i1\|CDS1 AT5G35630.3 AT5G35630.3\|GS2,GLN2,ATGSL1\|glutaminesynthetase2 | 8.25844818 |
| TRINITY_DN65341_c0_g1_i6\|CDS1,AT2G24530.1,AT2G24530.1 \| unknown protein | 8.24375738 |
| TRINITY_DN81821_c0_g1_i1\|CDS1,AT5G59240.1,AT5G59240.1 \| Ribosomal protein S8e family protein | 8.21083382 |

| | |
|---|---|
| TRINITY_DN109012_c0_g1_i1\|CDS1 AT5G60620.1 AT5G60620.1\|GPAT9\|glycerol-3-phosphateacyltransferase9 | 8.18328924 |
| TRINITY_DN98455_c0_g1_i1\|CDS1,AT1G30580.1,AT1G30580.1 \| GTP binding | 8.16097102 |
| TRINITY_DN107629_c0_g1_i1\|CDS1 AT2G26140.1 AT2G26140.1\|ftsh4\|FTSHprotease4 | 8.14376333 |
| TRINITY_DN51812_c0_g1_i1\|CDS1,AT1G20850.1,AT1G20850.1 \| XCP2 \| xylem cysteine peptidase 2 | 8.1293124 |
| TRINITY_DN52116_c0_g1_i1\|CDS1,AT1G69410.1,"AT1G69410.1 \| ATELF5A-3, ELF5A-3 \| eukaryotic elongation factor 5A-3 | 8.12579316 |
| TRINITY_DN53663_c0_g1_i4\|CDS1,AT3G55580.1,AT3G55580.1 \| Regulator of chromosome condensation (RCC1) family protein | 8.12000621 |
| TRINITY_DN99518_c0_g1_i1\|CDS1,AT2G15230.1,"AT2G15230.1 \| ATLIP1, LIP1 \| lipase 1 | 8.11169696 |
| TRINITY_DN51835_c0_g2_i1\|CDS1,AT4G19640.1,"AT4G19640.1 \| ARA7, ARA-7, ATRABF2B, ATRAB5B, RABF2B, ATRAB-F2B, RAB-F2B \| Ras-related small GTP-binding family protein | 8.10813496 |
| TRINITY_DN63627_c0_g1_i3\|CDS1,AT5G27690.1,AT5G27690.1 \| Heavy metal transport/detoxification superfamily protein | 8.07778269 |
| TRINITY_DN81973_c0_g1_i1\|CDS1,AT1G43890.3,"AT1G43890.3 \| ATRAB18, ATRABC1, RAB18-1, RABC1, ATRAB-C1, RAB18 \| RAB GTPASE HOMOLOG B18 | 8.06505771 |
| TRINITY_DN38934_c0_g1_i1\|CDS1 AT5G02610.1 AT5G02610.1\|RibosomalL29familyprotein | 8.05937707 |
| TRINITY_DN58284_c0_g1_i1\|CDS1,AT1G09640.1,"AT1G09640.1 \| Translation elongation factor EF1B, gamma chain | 8.05495265 |
| TRINITY_DN46342_c0_g1_i1\|CDS1 AT2G18110.1 AT2G18110.1\|TranslationelongationfactorEF1B/ribosomalproteinS6familyprotein | 8.0440843 |
| TRINITY_DN75320_c0_g1_i1\|CDS1,AT5G13710.2,"AT5G13710.2 \| SMT1, CPH \| sterol methyltransferase 1 | 8.02134082 |
| TRINITY_DN96851_c0_g1_i1\|CDS1,AT4G36130.1,AT4G36130.1 \| Ribosomal protein L2 family | 8.01266826 |
| TRINITY_DN37654_c0_g2_i2\|CDS1 AT2G37190.1 AT2G37190.1\|RibosomalproteinL11familyprotein | 8.01034134 |
| TRINITY_DN18984_c0_g2_i1\|CDS1 AT5G02960.1 AT5G02960.1\|RibosomalproteinS12/S23familyprotein | 7.99173126 |
| TRINITY_DN51971_c0_g1_i1\|CDS1,AT5G60390.3,AT5G60390.3 \| GTP binding Elongation factor Tu family protein | 7.9825199 |
| TRINITY_DN85475_c0_g1_i1\|CDS1,AT3G05930.1,AT3G05930.1 \| GLP8 \| germin-like protein 8 | 7.95753012 |
| TRINITY_DN63983_c0_g1_i1\|CDS1,AT1G51340.2,AT1G51340.2 \| MATE efflux family protein | 7.94245568 |
| TRINITY_DN8201_c0_g1_i1\|CDS1,AT3G23610.1,AT3G23610.1 \| DSPTP1 \| dual specificity protein phosphatase 1 | 7.94020462 |
| TRINITY_DN91632_c0_g1_i1\|CDS1,AT1G12440.2,AT1G12440.2 \| A20/AN1-like zinc finger family protein | 7.90058249 |
| TRINITY_DN68421_c0_g1_i1\|CDS1,AT1G01100.3,AT1G01100.3 \| 60S acidic ribosomal protein family | 7.89963655 |
| TRINITY_DN75198_c0_g1_i1\|CDS1,AT3G16640.1,AT3G16640.1 \| TCTP \| translationally controlled tumor protein | 7.89830074 |
| TRINITY_DN106810_c0_g1_i1\|CDS1 AT2G34480.1 AT2G34480.1\|RibosomalproteinL18ae/Lxfamilyprotein | 7.88832885 |

| | |
|---|---|
| TRINITY_DN4367_c0_g2_i1\|CDS1 AT2G09990.1 AT2G09990.1\|RibosomalproteinS5domain2-likesuperfamilyprotein | 7.87199375 |
| TRINITY_DN51640_c0_g1_i1\|CDS1,AT4G28680.4,AT4G28680.4 \| TYRDC \| L-tyrosine decarboxylase | 7.86659998 |
| TRINITY_DN47263_c0_g1_i1\|CDS1 AT5G58420.1 AT5G58420.1\|RibosomalproteinS4(RPS4A)familyprotein | 7.85656407 |
| TRINITY_DN1298_c0_g2_i1\|CDS1 AT1G09690.1 AT1G09690.1\|TranslationproteinSH3-likefamilyprotein | 7.82641534 |
| TRINITY_DN67703_c0_g1_i1\|CDS1,AT4G27270.1,AT4G27270.1 \| Quinone reductase family protein | 7.8032618 |
| TRINITY_DN65465_c0_g1_i1\|CDS1,AT5G58040.1,"AT5G58040.1 \| ATFIP1[V], FIPS5, ATFIPS5, FIP1[V] \| homolog of yeast FIP1 [V] | 7.75364305 |
| TRINITY_DN82099_c0_g1_i1\|CDS1,AT2G46210.1,AT2G46210.1 \| Fatty acid/sphingolipid desaturase | 7.72998843 |
| TRINITY_DN76504_c0_g1_i1\|CDS1,AT2G04520.1,"AT2G04520.1 \| Nucleic acid-binding, OB-fold-like protein | 7.7171101 |
| TRINITY_DN49790_c0_g1_i1\|CDS1,AT3G49010.4,"AT3G49010.4 \| ATBBC1, BBC1, RSU2 \| breast basic conserved 1 | 7.69171114 |
| TRINITY_DN90914_c0_g1_i1\|CDS1,AT4G14880.4,AT4G14880.4 \| OASA1 \| O-acetylserine (thiol) lyase (OAS-TL) isoform A1 | 7.69171114 |
| TRINITY_DN83506_c0_g1_i1\|CDS1,AT5G43340.1,"AT5G43340.1 \| PHT6, PHT1;6 \| phosphate transporter 1;6 | 7.68868981 |
| TRINITY_DN65777_c0_g1_i4\|CDS1,AT1G66330.2,AT1G66330.2 \| senescence-associated family protein | 7.66652843 |
| TRINITY_DN82693_c0_g1_i1\|CDS1,AT3G06680.1,AT3G06680.1 \| Ribosomal L29e protein family | 7.65772479 |
| TRINITY_DN5558_c0_g2_i1\|CDS1,AT1G67430.1,AT1G67430.1 \| Ribosomal protein L22p/L17e family protein | 7.64690872 |
| TRINITY_DN66114_c0_g1_i4\|CDS1,AT2G25530.1,AT2G25530.1 \| AFG1-like ATPase family protein | 7.63391592 |
| TRINITY_DN68696_c0_g1_i1\|CDS1,AT1G30230.1,"AT1G30230.1 \| Glutathione S-transferase, C-terminal-like;Translation elongation factor EF1B/ribosomal protein S6 | 7.62690976 |
| TRINITY_DN54452_c0_g1_i2\|CDS1,AT3G49340.1,AT3G49340.1 \| Cysteine proteinases superfamily protein | 7.59593159 |
| TRINITY_DN36241_c0_g2_i1\|CDS1 AT1G73230.1 AT1G73230.1\|Nascentpolypeptide-associatedcomplexNAC | 7.58795463 |
| TRINITY_DN21485_c0_g1_i1\|CDS1 AT5G23660.1 AT5G23660.1\|MTN3,SWEET12,AtSWEET12\|homologofMedicagotruncatulaMTN3 | 7.57605032 |
| TRINITY_DN65162_c0_g1_i8\|CDS1,AT3G45100.2,AT3G45100.2 \| SETH2 \| UDP-Glycosyltransferase superfamily protein | 7.54091469 |
| TRINITY_DN89186_c0_g1_i1\|CDS1,AT3G52930.1,AT3G52930.1 \| Aldolase superfamily protein | 7.52660261 |
| TRINITY_DN41153_c0_g3_i1\|CDS1 AT1G66580.1 AT1G66580.1\|SAG24,RPL10C\|senescenceassociatedgene24 | 7.51544903 |
| TRINITY_DN75916_c0_g1_i1\|CDS1,AT2G19770.1,AT2G19770.1 \| PRF5 \| profilin 5 | 7.50275678 |
| TRINITY_DN35805_c0_g1_i1\|CDS1 AT1G75630.1 AT1G75630.1\|AVA-P4\|vacuolarH+-pumpingATPase16kDaproteolipidsubunit4 | 7.49829739 |
| TRINITY_DN57594_c0_g1_i7\|CDS1,AT5G10100.2,AT5G10100.2 \| TPPI \| Haloacid dehalogenase-like hydrolase (HAD) superfamily protein | 7.49813637 |
| TRINITY_DN64779_c0_g1_i3\|CDS1,AT2G03340.1,AT2G03340.1 \| WRKY3 \| WRKY DNA-binding protein 3 | 7.49454227 |

| | |
|---|---|
| TRINITY_DN63617_c0_g1_i5\|CDS1,AT4G32180.3,"AT4G32180.3 \| ATPANK2, PANK2 \| pantothenate kinase 2 | 7.47543952 |
| TRINITY_DN66085_c0_g1_i2\|CDS1,AT3G43610.1,AT3G43610.1 \| Spc97 / Spc98 family of spindle pole body (SBP) component | 7.46824631 |
| TRINITY_DN56961_c0_g1_i1\|CDS1,AT5G58620.1,AT5G58620.1 \| zinc finger (CCCH-type) family protein | 7.45159563 |

**Supplemental Table 2.** List of the 100 unigenes that present a greater increase in their expression profiles in the vanilla root, at 2 dpi, in response to the *Fusarium* infection.

### 3) Top 100 list of transcripts that were repressed (down-regulated) at 2 dpi in the vanilla transcriptome, in response to *Fusarium*.

| **Name** | logFC |
|---|---|
| TRINITY_DN13643_c0_g1_i1\|CDS1 AT5G49690.1 AT5G49690.1\|UDP-Glycosyltransferasesuperfamilyprotein | -9.92521082 |
| TRINITY_DN65246_c0_g1_i6\|CDS1,AT1G70440.1,AT1G70440.1 \| SRO3 \| similar to RCD one 3 | -9.88868106 |
| TRINITY_DN63876_c0_g1_i8\|CDS1,AT3G07970.1,AT3G07970.1 \| QRT2 \| Pectin lyase-like superfamily protein | -9.88250824 |
| TRINITY_DN64387_c1_g1_i2\|CDS1,AT5G17420.1,"AT5G17420.1 \| IRX3, CESA7, ATCESA7, MUR10 \| Cellulose synthase family protein | -9.80596022 |
| TRINITY_DN63863_c0_g1_i2\|CDS1,AT5G18830.2,AT5G18830.2 \| SPL7 \| squamosa promoter binding protein-like 7 | -9.77861386 |
| TRINITY_DN66315_c0_g2_i1\|CDS1,AT5G04930.1,AT5G04930.1 \| ALA1 \| aminophospholipid ATPase 1 | -9.59414028 |
| TRINITY_DN64347_c0_g1_i5\|CDS1,AT1G76160.1,AT1G76160.1 \| sks5 \| SKU5 similar 5 | -9.43975058 |
| TRINITY_DN66332_c0_g2_i8\|CDS1,AT5G45140.1,AT5G45140.1 \| NRPC2 \| nuclear RNA polymerase C2 | -9.33315086 |
| TRINITY_DN66187_c0_g6_i10\|CDS1,AT4G05420.2,AT4G05420.2 \| DDB1A \| damaged DNA binding protein 1A | -9.32974378 |
| TRINITY_DN55988_c0_g1_i1\|CDS1,AT1G28520.2,AT1G28520.2 \| VOZ1 \| vascular plant one zinc finger protein | -9.09867864 |
| TRINITY_DN63540_c0_g1_i11\|CDS1,AT4G01040.1,AT4G01040.1 \| Glycosyl hydrolase superfamily protein | -9.0378224 |
| TRINITY_DN64517_c0_g2_i1\|CDS1,AT5G11240.1,AT5G11240.1 \| transducin family protein / WD-40 repeat family protein | -8.93951158 |
| TRINITY_DN58993_c0_g1_i2\|CDS1,AT3G02645.1,AT3G02645.1 \| Plant protein of unknown function (DUF247) | -8.813758 |
| TRINITY_DN59124_c0_g1_i3\|CDS1,AT4G28250.1,"AT4G28250.1 \| ATEXPB3, EXPB3, ATHEXP BETA 1.6 \| expansin B3 | -8.69571042 |
| TRINITY_DN64290_c0_g1_i9\|CDS1,AT2G47180.1,"AT2G47180.1 \| AtGolS1, GolS1 \| galactinol synthase 1 | -8.5633944 |

| | |
|---|---|
| TRINITY_DN64387_c1_g1_i8\|CDS1,AT5G17420.1,"AT5G17420.1 \| IRX3, CESA7, ATCESA7, MUR10 \| Cellulose synthase family protein | -8.47130866 |
| TRINITY_DN58334_c0_g1_i5\|CDS1,AT1G31600.2,AT1G31600.2 \| RNA-binding (RRM/RBD/RNP motifs) family protein | -8.28543786 |
| TRINITY_DN54718_c0_g1_i2\|CDS1,AT5G01230.1,AT5G01230.1 \| S-adenosyl-L-methionine-dependent methyltransferases superfamily protein | -8.2406168 |
| TRINITY_DN63940_c0_g1_i3\|CDS1,AT1G48100.1,AT1G48100.1 \| Pectin lyase-like superfamily protein | -8.22359788 |
| TRINITY_DN65451_c0_g1_i1\|CDS1,AT1G13030.1,AT1G13030.1 \| sphere organelles protein-related | -8.10617133 |
| TRINITY_DN64510_c0_g1_i1\|CDS1,AT5G52060.1,"AT5G52060.1 \| ATBAG1, BAG1 \| BCL-2-associated athanogene 1 | -8.09979924 |
| TRINITY_DN52123_c0_g1_i2\|CDS1,AT4G11120.1,"AT4G11120.1 \| translation elongation factor Ts (EF-Ts), putative | -7.98755131 |
| TRINITY_DN58067_c0_g2_i11\|CDS1,AT1G28510.1,AT1G28510.1 \| Optic atrophy 3 protein (OPA3) | -7.97518145 |
| TRINITY_DN53548_c0_g1_i1\|CDS1,AT1G57560.1,"AT1G57560.1 \| AtMYB50, MYB50 \| myb domain protein 50 | -7.80836399 |
| TRINITY_DN65632_c1_g2_i4\|CDS1,AT3G04680.2,AT3G04680.2 \| CLPS3 \| CLP-similar protein 3 | -7.78693227 |
| TRINITY_DN44034_c0_g1_i1\|CDS1 AT4G22000.1 AT4G22000.1\|unknownprotein | -7.785356 |
| TRINITY_DN48812_c0_g1_i3\|CDS1,AT2G20980.1,AT2G20980.1 \| MCM10 \| minichromosome maintenance 10 | -7.76051928 |
| TRINITY_DN65911_c0_g1_i5\|CDS1,AT1G75450.2,AT1G75450.2 \| CKX5 \| cytokinin oxidase 5 | -7.75951671 |
| TRINITY_DN63189_c0_g1_i3\|CDS1,AT1G22170.1,AT1G22170.1 \| Phosphoglycerate mutase family protein | -7.74632216 |
| TRINITY_DN57452_c0_g1_i2\|CDS1,AT4G29810.1,"AT4G29810.1 \| ATMKK2, MKK2, MK1 \| MAP kinase kinase 2 | -7.74151473 |
| TRINITY_DN65018_c0_g1_i5\|CDS1,AT3G25500.1,"AT3G25500.1 \| AFH1, FH1, AHF1, ATFH1 \| formin homology 1 " | -7.73504914 |
| TRINITY_DN66314_c0_g1_i1\|CDS1,AT4G23540.1,AT4G23540.1 \| ARM repeat superfamily protein | -7.70833118 |
| TRINITY_DN64154_c0_g1_i4\|CDS1,AT3G07670.1,AT3G07670.1 \| Rubisco methyltransferase family protein | -7.68762909 |
| TRINITY_DN64774_c0_g1_i5\|CDS1,AT1G03620.1,AT1G03620.1 \| ELMO/CED-12 family protein | -7.6839425 |
| TRINITY_DN46681_c0_g1_i2\|CDS1 AT1G62340.1 AT1G62340.1\|ALE1,ALE\|PA-domaincontainingsubtilasefamilyprotein | -7.64223606 |
| TRINITY_DN63983_c0_g1_i2\|CDS1,AT1G51340.2,AT1G51340.2 \| MATE efflux family protein | -7.63504655 |
| TRINITY_DN43149_c0_g1_i2\|CDS1 AT5G19740.1 AT5G19740.1\|PeptidaseM28familyprotein | -7.62104217 |
| TRINITY_DN65829_c1_g1_i7\|CDS1,AT4G21390.1,AT4G21390.1 \| B120 \| S-locus lectin protein kinase family protein | -7.6133339 |
| TRINITY_DN58817_c0_g1_i2\|CDS1,AT3G15070.2,AT3G15070.2 \| RING/U-box superfamily protein | -7.58092057 |
| TRINITY_DN57034_c0_g1_i1\|CDS1,AT1G63100.1,AT1G63100.1 \| GRAS family transcription factor | -7.57700984 |
| TRINITY_DN63795_c0_g1_i4\|CDS1,AT4G38050.1,AT4G38050.1 \| Xanthine/uracil permease family protein | -7.55088594 |

| | |
|---|---|
| TRINITY_DN64156_c0_g1_i2\|CDS1,AT1G32740.1,AT1G32740.1 \| SBP (S-ribonuclease binding protein) family protein | -7.5184247 |
| TRINITY_DN58092_c0_g2_i1\|CDS1,AT5G66750.1,"AT5G66750.1 \| DDM1, CHR01, CHR1, CHA1, SOM4, SOM1, ATDDM1 \| chromatin remodeling 1 | -7.51541208 |
| TRINITY_DN57211_c0_g1_i2\|CDS1,AT1G54820.1,AT1G54820.1 \| Protein kinase superfamily protein | -7.46318928 |
| TRINITY_DN57314_c0_g1_i1\|CDS1,AT3G05675.1,AT3G05675.1 \| BTB/POZ domain-containing protein | -7.4254775 |
| TRINITY_DN53143_c0_g1_i1\|CDS1,AT2G26170.2,"AT2G26170.2 \| CYP711A1 \| cytochrome P450, family 711, subfamily A, polypeptide 1 | -7.42238379 |
| TRINITY_DN63476_c0_g3_i8\|CDS1,AT3G02690.1,AT3G02690.1 \| nodulin MtN21 /EamA-like transporter family protein | -7.39379106 |
| TRINITY_DN57157_c0_g1_i1\|CDS1,AT1G75500.2,AT1G75500.2 \| WAT1 \| Walls Are Thin 1 | -7.36720836 |
| TRINITY_DN64542_c0_g1_i2\|CDS1,AT4G19191.1,AT4G19191.1 \| Tetratricopeptide repeat (TPR)-like superfamily protein | -7.36700928 |
| TRINITY_DN47476_c0_g1_i3\|CDS1 AT3G05800.1 AT3G05800.1\|AIF1\|AtBS1(activation-taggedBRI1suppressor1)-interactingfactor1 | -7.36630757 |
| TRINITY_DN64881_c0_g1_i3\|CDS1,AT4G32050.1,AT4G32050.1 \| neurochondrin family protein | -7.3074634 |
| TRINITY_DN63346_c0_g1_i9\|CDS1,AT1G79620.1,AT1G79620.1 \| Leucine-rich repeat protein kinase family protein | -7.29801505 |
| TRINITY_DN63679_c0_g1_i2\|CDS1,AT1G76520.2,AT1G76520.2 \| Auxin efflux carrier family protein | -7.25808457 |
| TRINITY_DN63804_c0_g1_i1\|CDS1,AT4G05150.1,AT4G05150.1 \| Octicosapeptide/Phox/Bem1p family protein | -7.22982297 |
| TRINITY_DN64556_c0_g1_i1\|CDS1,AT3G18670.1,AT3G18670.1 \| Ankyrin repeat family protein | -7.20624074 |
| TRINITY_DN66105_c0_g1_i8\|CDS1,AT3G02840.1,AT3G02840.1 \| ARM repeat superfamily protein | -7.10028842 |
| TRINITY_DN55222_c0_g1_i3\|CDS1,AT3G55820.1,AT3G55820.1 \| Fasciclin-like arabinogalactan family protein | -7.07215605 |
| TRINITY_DN57539_c0_g1_i4\|CDS1,AT5G38520.1,AT5G38520.1 \| alpha/beta-Hydrolases superfamily protein | -7.03720202 |
| TRINITY_DN64632_c0_g1_i8\|CDS1,AT5G14620.1,"AT5G14620.1 \| DRM2, DMT7 \| domains rearranged methyltransferase 2 | -7.00383882 |
| TRINITY_DN66035_c2_g1_i2\|CDS1,AT1G63430.1,AT1G63430.1 \| Leucine-rich repeat protein kinase family protein | -6.97312931 |
| TRINITY_DN55366_c0_g1_i5\|CDS1,AT1G68730.1,AT1G68730.1 \| Zim17-type zinc finger protein | -6.9382651 |
| TRINITY_DN64295_c0_g1_i4\|CDS1,AT1G68930.1,AT1G68930.1 \| pentatricopeptide (PPR) repeat-containing protein | -6.93803321 |
| TRINITY_DN55080_c0_g2_i3\|CDS1,AT3G49940.1,AT3G49940.1 \| LBD38 \| LOB domain-containing protein 38 | -6.89317959 |
| TRINITY_DN65714_c2_g1_i2\|CDS1,AT3G12670.1,AT3G12670.1 \| emb2742 \| CTP synthase family protein | -6.8854109 |
| TRINITY_DN47612_c0_g1_i3\|CDS1 AT1G52740.1 AT1G52740.1\|HTA9\|histoneH2Aprotein9 | -6.86373028 |

| | |
|---|---|
| TRINITY_DN58768_c0_g1_i2\|CDS1,AT1G48570.1,AT1G48570.1 \| zinc finger (Ran-binding) family protein | -6.84300117 |
| TRINITY_DN66491_c0_g1_i7\|CDS1,AT1G11330.1,AT1G11330.1 \| S-locus lectin protein kinase family protein \| | -6.83484292 |
| TRINITY_DN55954_c0_g1_i2\|CDS1,AT3G07080.1,AT3G07080.1 \| EamA-like transporter family | -6.80888264 |
| TRINITY_DN44161_c0_g1_i1\|CDS1 AT4G01130.1 AT4G01130.1\|GDSL-likeLipase/Acylhydrolasesuperfamilyprotein | -6.78831327 |
| TRINITY_DN65833_c0_g1_i2\|CDS1,AT1G67930.1,AT1G67930.1 \| Golgi transport complex protein-related | -6.77327405 |
| TRINITY_DN63489_c0_g1_i6\|CDS1,AT1G01140.1,"AT1G01140.1 \| CIPK9, SnRK3.12, PKS6 \| CBL-interacting protein kinase 9 | -6.73676847 |
| TRINITY_DN66017_c0_g1_i1\|CDS1,AT1G63300.1,AT1G63300.1 \| Myosin heavy chain-related protein | -6.73064646 |
| TRINITY_DN63167_c0_g1_i2\|CDS1,AT4G21070.1,"AT4G21070.1 \| ATBRCA1, BRCA1 \| breast cancer susceptibility1 | -6.72676948 |
| TRINITY_DN65946_c1_g1_i6\|CDS1,AT5G13050.1,AT5G13050.1 \| 5-FCL \| 5-formyltetrahydrofolate cycloligase | -6.62490521 |
| TRINITY_DN53480_c0_g2_i7\|CDS1,AT2G30933.2,AT2G30933.2 \| Carbohydrate-binding X8 domain superfamily protein | -6.61942426 |
| TRINITY_DN65714_c2_g1_i8\|CDS1,AT3G12670.1,AT3G12670.1 \| emb2742 \| CTP synthase family protein | -6.61162669 |
| TRINITY_DN52065_c0_g1_i1\|CDS1,AT1G79620.1,AT1G79620.1 \| Leucine-rich repeat protein kinase family protein | -6.58287339 |
| TRINITY_DN57690_c0_g1_i4\|CDS1,AT5G46860.1,"AT5G46860.1 \| VAM3, ATVAM3, SYP22, ATSYP22, SGR3 \| Syntaxin/t-SNARE family protein | -6.58159722 |
| TRINITY_DN50595_c0_g2_i1\|CDS1,AT5G65640.2,AT5G65640.2 \| bHLH093 \| beta HLH protein 93 | -6.54859264 |
| TRINITY_DN63991_c0_g1_i3\|CDS1,AT1G12420.1,AT1G12420.1 \| ACR8 \| ACT domain repeat 8 | -6.54185398 |
| TRINITY_DN66122_c0_g1_i5\|CDS1,AT5G03760.1,"AT5G03760.1 \| ATCSLA09, CSLA09, ATCSLA9, CSLA9, RAT4 \| Nucleotide-diphospho-sugar transferases superfamily protein | -6.53446643 |
| TRINITY_DN65353_c0_g2_i3\|CDS1,AT4G30980.1,AT4G30980.1 \| LRL2 \| LJRHL1-like 2 | -6.51289691 |
| TRINITY_DN56491_c0_g1_i5\|CDS1,AT3G15140.1,"AT3G15140.1 \| Polynucleotidyl transferase, ribonuclease H-like superfamily protein | -6.5048057 |
| TRINITY_DN55871_c0_g1_i2\|CDS1,AT2G25950.1,AT2G25950.1 \| Protein of unknown function (DUF1000) | -6.4651861 |
| TRINITY_DN54413_c0_g1_i3\|CDS1,AT5G66650.1,AT5G66650.1 \| Protein of unknown function (DUF607) | -6.38764347 |
| TRINITY_DN54413_c0_g1_i3\|CDS1,AT5G66650.1,AT5G66650.1 \| Protein of unknown function (DUF607) | -6.38764347 |
| TRINITY_DN65776_c0_g1_i8\|CDS1,AT3G55080.1,AT3G55080.1 \| SET domain-containing protein | -6.35014647 |
| TRINITY_DN97652_c0_g1_i1\|CDS1,AT1G33920.1,"AT1G33920.1 \| ATPP2-A4, PP2-A4 \| phloem protein 2-A4 | -6.33689943 |
| TRINITY_DN64275_c0_g1_i3\|CDS1,AT1G76730.1,AT1G76730.1 \| NagB/RpiA/CoA transferase-like superfamily protein | -6.31352484 |

| | |
|---|---|
| TRINITY_DN63785_c0_g1_i4\|CDS1,AT1G10830.2,"AT1G10830.2 \| Z-ISO, Z-ISO1.2 \| 15-cis-zeta-carotene isomerase | -6.30235002 |
| TRINITY_DN64329_c0_g1_i7\|CDS1,AT3G62080.1,AT3G62080.1 \| SNF7 family protein | -6.30155614 |
| TRINITY_DN64426_c0_g1_i4\|CDS1,AT4G21470.1,"AT4G21470.1 \| ATFMN/FHY, FMN/FHY \| riboflavin kinase/FMN hydrolase | -6.29981707 |
| TRINITY_DN50893_c0_g1_i1\|CDS1,AT1G48405.1,AT1G48405.1 \| Kinase interacting (KIP1-like) family protein | -6.29897971 |
| TRINITY_DN54791_c0_g1_i4\|CDS1,AT2G02500.1,"AT2G02500.1 \| ISPD, ATMEPCT, MCT \| Nucleotide-diphospho-sugar transferases superfamily protein | -6.24989627 |
| TRINITY_DN55554_c0_g1_i3\|CDS1,AT3G12300.1,AT3G12300.1 \| unknown protein | -6.23302622 |
| TRINITY_DN64249_c2_g2_i5\|CDS1,AT5G23960.1,"AT5G23960.1 \| ATTPS21, TPS21 \| terpene synthase 21 | -6.15381606 |
| TRINITY_DN53405_c0_g1_i1\|CDS1,AT1G11260.1,"AT1G11260.1 \| STP1, ATSTP1 \| sugar transporter 1 | -5.96183693 |
| TRINITY_DN46469_c0_g1_i2\|CDS1 AT5G60920.1 AT5G60920.1\|COB\|COBRA-likeextracellularglycosyl-phosphatidylinositol-anchoredproteinfamily | -5.93156636 |
| TRINITY_DN56596_c0_g1_i1\|CDS1,AT1G09000.1,"AT1G09000.1 \| ANP1, MAPKKK1, NP1 \| NPK1-related protein kinase 1 | -5.7895658 |

**Supplemental Table 3**. List of the 100 unigenes that present a greater decrease in their expression profiles in the vanilla root, at 2 dpi, in response to the *Fusarium* infection.

4) **Main categories of gene ontology corresponding to the transcripts of the late response (10 dpi) of vanilla before *Fusarium*.**

| Name | elements | p-value |
|---|---|---|
| C1-metabolism | 1 | 0.12557647 |
| lipid metabolism | 3 | 0.15029475 |
| minor CHO metabolism | 1 | 0.15737607 |
| amino acid metabolism | 1 | 0.17168173 |
| secondary metabolism | 6 | 0.23233808 |
| misc | 9 | 0.38287291 |
| hormone metabolism | 5 | 0.4045416 |
| signalling | 8 | 0.41724757 |
| cell | 6 | 0.53095537 |
| nucleotide metabolism | 2 | 0.65165411 |
| RNA | 27 | 0.87281801 |
| DNA | 3 | 0.88021311 |
| development | 4 | 0.88639234 |
| Biodegradation of Xenobiotics | 1 | 0.88755777 |
| Co-factor and vitamine metabolism | 2 | 0.92008579 |
| not assigned | 32 | 0.92147782 |
| transport | 9 | 0.92276036 |

| | | |
|---|---|---|
| protein | 20 | 0.92797027 |
| stress | 5 | 0.95738779 |
| major CHO metabolism | 2 | 0.95999263 |
| cell wall | 2 | 1 |
| signalling.G-proteins | 2 | 0.0397174 |
| protein.postranslational modification.kinase | 2 | 0.07359213 |
| protein.degradation.ubiquitin.E3 | 2 | 0.08502295 |
| protein.degradation.metalloprotease | 1 | 0.08537648 |
| RNA.processing | 1 | 0.09902265 |
| RNA.processing.RNA helicase | 1 | 0.09902265 |
| major CHO metabolism.degradation.starch | 1 | 0.09902265 |
| major CHO metabolism.degradation.starch.starch cleavage | 1 | 0.09902265 |
| signalling.phosphorelay | 1 | 0.10394019 |
| protein.postranslational modification.kinase.receptor like cytoplasmatic kinase VI | 1 | 0.10394019 |
| major CHO metabolism.degradation.sucrose | 1 | 0.11435606 |
| major CHO metabolism.degradation.sucrose.fructokinase | 1 | 0.11435606 |
| protein.degradation.ubiquitin.E3.RING | 1 | 0.11435606 |
| cell.vesicle transport | 2 | 0.13670767 |
| lipid metabolism.Phospholipid synthesis | 1 | 0.13763674 |
| RNA.regulation of transcription.C3H zinc finger family | 1 | 0.15057097 |
| protein.degradation.aspartate protease | 1 | 0.15057097 |
| minor CHO metabolism.others | 1 | 0.15737607 |
| protein.postranslational modification | 8 | 0.16139442 |
| cell.organisation | 3 | 0.16229591 |

| | | |
|---|---|---|
| amino acid metabolism.synthesis | 1 | 0.17168173 |
| amino acid metabolism.synthesis.aspartate family | 1 | 0.17168173 |
| amino acid metabolism.synthesis.aspartate family.lysine | 1 | 0.17168173 |
| amino acid metabolism.synthesis.aspartate family.lysine.LL-diaminopimelic acid aminotransferase | 1 | 0.17168173 |
| signalling.in sugar and nutrient physiology | 1 | 0.17918969 |
| signalling.in sugar and nutrient physiology | 1 | 0.17918969 |
| protein.synthesis.ribosomal protein.eukaryotic.60S subunit.L27 | 1 | 0.18693911 |
| protein.degradation.ubiquitin.ubiquitin protease | 1 | 0.18693911 |
| protein.synthesis | 2 | 0.19215396 |
| protein.synthesis.ribosomal protein | 2 | 0.19215396 |
| protein.synthesis.ribosomal protein.eukaryotic | 2 | 0.19215396 |
| protein.synthesis.ribosomal protein.eukaryotic.60S subunit | 2 | 0.19215396 |
| misc.peroxidases | 2 | 0.21595725 |
| Co-factor and vitamine metabolism.biotin | 1 | 0.22041519 |
| cell.division | 1 | 0.22941817 |
| RNA.regulation of transcription.WRKY domain transcription factor family | 5 | 0.23135548 |
| cell wall.degradation | 1 | 0.23867977 |
| cell wall.degradation.pectate lyases and polygalacturonases | 1 | 0.23867977 |
| cell wall.modification | 1 | 0.23867977 |
| transport.metal | 1 | 0.24820213 |
| hormone metabolism.brassinosteroid | 1 | 0.24820213 |
| hormone metabolism.brassinosteroid.synthesis-degradation | 1 | 0.24820213 |
| hormone metabolism.brassinosteroid.synthesis-degradation.BRs | 1 | 0.24820213 |
| hormone metabolism.brassinosteroid.synthesis-degradation.BRs.DWF4 | 1 | 0.24820213 |

| | | |
|---|---|---|
| DNA.unspecified | 1 | 0.26803653 |
| Co-factor and vitamine metabolism.riboflavin | 1 | 0.27835164 |
| Co-factor and vitamine metabolism.riboflavin.GTP cyclohydrolase II | 1 | 0.27835164 |
| hormone metabolism.gibberelin | 1 | 0.3222884 |
| hormone metabolism.gibberelin.induced-regulated-responsive-activated | 1 | 0.3222884 |
| RNA.RNA binding | 1 | 0.33394393 |
| secondary metabolism.simple phenols | 1 | 0.34586789 |
| protein.degradation | 8 | 0.34758147 |
| misc.glutathione S transferases | 1 | 0.37051839 |
| protein.postranslational modification.kinase.receptor like cytoplasmatic kinase VII | 1 | 0.37051839 |
| protein.degradation.ubiquitin.E3.BTB/POZ Cullin3 | 1 | 0.39623136 |
| protein.degradation.ubiquitin.E3.BTB/POZ Cullin3.BTB/POZ | 1 | 0.39623136 |
| nucleotide metabolism.phosphotransfer and pyrophosphatases | 1 | 0.39623136 |
| nucleotide metabolism.phosphotransfer and pyrophosphatases.uridylate kinase | 1 | 0.39623136 |
| signalling.lipids | 1 | 0.4094822 |
| RNA.regulation of transcription.unclassified | 2 | 0.41260032 |
| signalling.calcium | 1 | 0.42299311 |
| not assigned.no ontology.pentatricopeptide (PPR) repeat-containing protein | 3 | 0.43486414 |
| RNA.regulation of transcription.NAC domain transcription factor family | 1 | 0.43676154 |
| secondary metabolism.isoprenoids | 1 | 0.45078462 |
| secondary metabolism.isoprenoids.terpenoids | 1 | 0.45078462 |
| misc.acid and other phosphatases | 1 | 0.46505911 |
| protein.targeting | 1 | 0.47958147 |
| protein.targeting.chloroplast | 1 | 0.47958147 |

| | | |
|---|---|---|
| protein.degradation.cysteine protease | 1 | 0.47958147 |
| lipid metabolism.FA synthesis and FA elongation | 2 | 0.48250481 |
| lipid metabolism.FA synthesis and FA elongation.acyl coa ligase | 2 | 0.48250481 |
| transport.metabolite transporters at the mitochondrial membrane | 1 | 0.4943478 |
| transport.ammonium | 1 | 0.52459506 |
| RNA.regulation of transcription.MYB domain transcription factor family | 4 | 0.54371804 |
| DNA.repair | 2 | 0.54720479 |
| misc.cytochrome P450 | 2 | 0.54720479 |
| not assigned.no ontology.AT hook motif-containing protein | 1 | 0.57167885 |
| transport.amino acids | 2 | 0.58109147 |
| protein.degradation.serine protease | 1 | 0.58780829 |
| protein.degradation.ubiquitin | 4 | 0.59212104 |
| not assigned.unknown | 15 | 0.59976659 |
| protein.synthesis.ribosomal protein.eukaryotic.60S subunit.L22 | 1 | 0.60414505 |
| secondary metabolism.phenylpropanoids | 2 | 0.60421463 |
| signalling.receptor kinases | 2 | 0.62774207 |
| RNA.regulation of transcription.HSF,Heat-shock transcription factor family | 3 | 0.65119895 |
| signalling.receptor kinases.thaumatin like | 1 | 0.65433263 |
| hormone metabolism.salicylic acid | 2 | 0.66374788 |
| hormone metabolism.salicylic acid.synthesis-degradation | 2 | 0.66374788 |
| RNA.regulation of transcription | 25 | 0.68377215 |
| not assigned.no ontology | 17 | 0.71176837 |
| secondary metabolism.phenylpropanoids.lignin biosynthesis | 1 | 0.72372267 |
| secondary metabolism.phenylpropanoids.lignin biosynthesis.PAL | 1 | 0.72372267 |

| | | |
|---|---|---|
| secondary metabolism.flavonoids | 2 | 0.75071409 |
| hormone metabolism.jasmonate | 1 | 0.75933344 |
| hormone metabolism.jasmonate.synthesis-degradation | 1 | 0.75933344 |
| hormone metabolism.jasmonate.synthesis-degradation.12-Oxo-PDA-reductase | 1 | 0.75933344 |
| RNA.regulation of transcription.Chromatin Remodeling Factors | 1 | 0.77733755 |
| RNA.regulation of transcription.PHOR1 | 1 | 0.77733755 |
| secondary metabolism.flavonoids.dihydroflavonols | 1 | 0.79546204 |
| protein.glycosylation | 1 | 0.81369758 |
| signalling.receptor kinases.DUF 26 | 1 | 0.81369758 |
| RNA.regulation of transcription.putative transcription regulator | 3 | 0.82649531 |
| nucleotide metabolism.deoxynucleotide metabolism | 1 | 0.83203467 |
| nucleotide metabolism.deoxynucleotide metabolism.ribonucleoside-diphosphate reductase | 1 | 0.83203467 |
| secondary metabolism.flavonoids.chalcones | 1 | 0.85046363 |
| secondary metabolism.flavonoids.chalcones.naringenin-chalcone synthase | 1 | 0.85046363 |
| RNA.regulation of transcription.HB,Homeobox transcription factor family | 2 | 0.85406808 |
| misc.beta 1,3 glucan hydrolases | 1 | 0.86897465 |
| misc.beta 1,3 glucan hydrolases.glucan endo-1,3-beta-glucosidase | 1 | 0.86897465 |
| misc.short chain dehydrogenase/reductase (SDR) | 1 | 0.86897465 |
| development.unspecified | 4 | 0.88639234 |
| RNA.regulation of transcription.C2H2 zinc finger family | 1 | 0.88755777 |
| transport.p- and v-ATPases | 1 | 0.9062029 |
| transport.ABC transporters and multidrug resistance systems | 3 | 0.92360046 |
| RNA.regulation of transcription.bHLH,Basic Helix-Loop-Helix family | 1 | 0.94363844 |
| stress.biotic | 5 | 0.95738779 |

| | | |
|---|---|---|
| major CHO metabolism.degradation | 2 | 0.95999263 |
| misc.GDSL-motif lipase | 1 | 0.9811989 |

**Supplemental Table 4.** List of the main functional categories, determined with the Mapman 3.0 software, to which the annotated transcripts belong, which show differential expression at 10 dpi, in the vanilla transcriptome in response to *Fusarium*.

5) **List of transcripts expressed (up-regulated), during the late response, at 10 dpi, in the vanilla transcriptome, in response to *Fusarium*.**

| Name | LogFC |
|---|---|
| TRINITY_DN64179_c0_g1_i1\|CDS1,AT3G22830.1,"AT3G22830.1 \| AT-HSFA6B, HSFA6B \| heat shock transcription factor A6B | 4.225012229 |
| TRINITY_DN56934_c0_g1_i4\|CDS1,AT3G50660.1,"AT3G50660.1 \| DWF4, CYP90B1, CLM, SNP2, SAV1, PSC1 \| Cytochrome P450 superfamily protein | 4.26837477 |
| TRINITY_DN56934_c0_g1_i4\|CDS1,AT3G50660.1,"AT3G50660.1 \| DWF4, CYP90B1, CLM, SNP2, SAV1, PSC1 \| Cytochrome P450 superfamily protein | 4.26837477 |
| TRINITY_DN63940_c0_g1_i2\|CDS1,AT1G48100.1,AT1G48100.1 \| Pectin lyase-like superfamily protein | 6.398395088 |
| TRINITY_DN63629_c0_g1_i1\|CDS1,AT5G58320.2,AT5G58320.2 \| Kinase interacting (KIP1-like) family protein | 6.528636349 |
| TRINITY_DN48106_c0_g1_i4\|CDS1,AT4G13730.1,AT4G13730.1 \| Ypt/Rab-GAP domain of gyp1p superfamily protein | 6.566074127 |
| TRINITY_DN48106_c0_g1_i4\|CDS1,AT4G13730.1,AT4G13730.1 \| Ypt/Rab-GAP domain of gyp1p superfamily protein | 6.566074127 |
| TRINITY_DN54908_c0_g1_i3\|CDS1,AT2G37195.1,AT2G37195.1 \| unknown protein | 6.733962876 |
| TRINITY_DN54908_c0_g1_i3\|CDS1,AT2G37195.1,AT2G37195.1 \| unknown protein | 6.733962876 |
| TRINITY_DN53703_c0_g1_i4\|CDS1,AT3G06880.2,AT3G06880.2 \| Transducin/WD40 repeat-like superfamily protein | 6.746130092 |

| | |
|---|---|
| TRINITY_DN53703_c0_g1_i4\|CDS1,AT3G06880.2,AT3G06880.2 \| Transducin/WD40 repeat-like superfamily protein | 6.746130092 |
| TRINITY_DN54601_c0_g1_i4\|CDS1,AT1G69560.1,"AT1G69560.1 \| MYB105, LOF2, ATMYB105 \| myb domain protein 105 | 6.765645558 |
| TRINITY_DN54601_c0_g1_i4\|CDS1,AT1G69560.1,"AT1G69560.1 \| MYB105, LOF2, ATMYB105 \| myb domain protein 105 | 6.765645558 |
| TRINITY_DN19411_c0_g1_i1\|CDS1 AT4G15000.1 AT4G15000.1\|RibosomalL27eproteinfamily | 6.794627907 |
| TRINITY_DN19411_c0_g1_i1\|CDS1 AT4G15000.1 AT4G15000.1\|RibosomalL27eproteinfamily | 6.794627907 |
| TRINITY_DN55353_c0_g1_i\|CDS1,AT1G68260.1,AT1G68260.1 \| Thioesterase superfamily protein | 6.875134674 |
| TRINITY_DN55353_c0_g1_i\|CDS1,AT1G68260.1,AT1G68260.1 \| Thioesterase superfamily protein | 6.875134674 |
| TRINITY_DN52471_c0_g1_i1\|CDS1,AT3G13870.2,AT3G13870.2 \| RHD3 \| Root hair defective 3 GTP-binding protein (RHD3) | 6.891933845 |
| TRINITY_DN52471_c0_g1_i1\|CDS1,AT3G13870.2,AT3G13870.2 \| RHD3 \| Root hair defective 3 GTP-binding protein (RHD3) | 6.891933845 |
| TRINITY_DN58147_c0_g1_i4\|CDS1,AT4G31150.1,AT4G31150.1 \| endonuclease V family protein | 6.936084415 |
| TRINITY_DN58147_c0_g1_i4\|CDS1,AT4G31150.1,AT4G31150.1 \| endonuclease V family protein | 6.936084415 |
| TRINITY_DN63629_c0_g1_i4\|CDS1,AT5G58320.2,AT5G58320.2 \| Kinase interacting (KIP1-like) family protein | 7.082914033 |
| TRINITY_DN64037_c0_g1_i8\|CDS1,AT1G04910.1,AT1G04910.1 \| O-fucosyltransferase family protein | 7.237653757 |
| TRINITY_DN63726_c0_g1_i1\|CDS1,AT2G41900.1,AT2G41900.1 \| CCCH-type zinc finger protein with ARM repeat domain | 7.283804333 |
| TRINITY_DN66777_c0_g6_i4\|CDS1,AT3G03800.1,"AT3G03800.1 \| SYP131, ATSYP131 \| syntaxin of plants 131 | 7.330551914 |
| TRINITY_DN64795_c0_g1_i3\|CDS1,AT1G31410.1,AT1G31410.1 \| putrescine-binding periplasmic protein-related | 7.407896764 |
| TRINITY_DN65617_c0_g1_i3\|CDS1,AT2G13440.1,AT2G13440.1 \| glucose-inhibited division family A protein | 7.473133446 |

| | |
|---|---|
| TRINITY_DN63515_c0_g1_i2\|CDS1,AT3G29230.1,AT3G29230.1 \| Tetratricopeptide repeat (TPR)-like superfamily protein | 7.494087292 |
| TRINITY_DN66242_c0_g1_i2\|CDS1,AT5G45720.2,AT5G45720.2 \| AAA-type ATPase family protein | 7.541842897 |
| TRINITY_DN52389_c0_g1_i3\|CDS1,AT5G03500.3,"AT5G03500.3 \| Mediator complex, subunit Med7 | 7.676455154 |
| TRINITY_DN52389_c0_g1_i3\|CDS1,AT5G03500.3,"AT5G03500.3 \| Mediator complex, subunit Med7 | 7.676455154 |
| TRINITY_DN48795_c0_g1_i1\|CDS1,AT1G19310.1,AT1G19310.1 \| RING/U-box superfamily protein | 7.856010574 |
| TRINITY_DN48795_c0_g1_i1\|CDS1,AT1G19310.1,AT1G19310.1 \| RING/U-box superfamily protein | 7.856010574 |
| TRINITY_DN65353_c0_g2_i2\|CDS1,AT4G30980.1,AT4G30980.1 \| LRL2 \| LJRHL1-like 2 | 8.17422348 |
| TRINITY_DN56473_c0_g1_i7\|CDS1,AT1G72650.2,AT1G72650.2 \| TRFL6 \| TRF-like 6 | 8.244885971 |
| TRINITY_DN56473_c0_g1_i7\|CDS1,AT1G72650.2,AT1G72650.2 \| TRFL6 \| TRF-like 6 | 8.244885971 |
| TRINITY_DN58049_c0_g1_i5\|CDS1,AT3G21510.1,AT3G21510.1 \| AHP1 \| histidine-containing phosphotransmitter 1 | 8.54748858 |
| TRINITY_DN58049_c0_g1_i5\|CDS1,AT3G21510.1,AT3G21510.1 \| AHP1 \| histidine-containing phosphotransmitter 1 | 8.54748858 |
| TRINITY_DN65306_c0_g1_i3\|CDS1,AT2G45880.1,"AT2G45880.1 \| BMY4, BAM7 \| beta-amylase 7 | 8.640296237 |
| TRINITY_DN65890_c0_g1_i9\|CDS1,AT3G45830.1,AT3G45830.1 \| unknown protein | 8.665890337 |
| TRINITY_DN58407_c0_g1_i1\|CDS1,AT1G31650.1,"AT1G31650.1 \| ATROPGEF14, ROPGEF14 \| RHO guanyl-nucleotide exchange factor 14 | 8.881266887 |
| TRINITY_DN58407_c0_g1_i1\|CDS1,AT1G31650.1,"AT1G31650.1 \| ATROPGEF14, ROPGEF14 \| RHO guanyl-nucleotide exchange factor 14 | 8.881266887 |
| TRINITY_DN58457_c0_g1_i5\|CDS1,AT1G07430.1,AT1G07430.1 \| HAI2 \| highly ABA-induced PP2C gene 2 | 9.381107811 |
| TRINITY_DN58457_c0_g1_i5\|CDS1,AT1G07430.1,AT1G07430.1 \| HAI2 \| highly ABA-induced PP2C gene 2 | 9.381107811 |
| TRINITY_DN59234_c0_g1_i2\|CDS1,AT4G23940.1,AT4G23940.1 \| FtsH extracellular protease family | 9.798896331 |
| TRINITY_DN59234_c0_g1_i2\|CDS1,AT4G23940.1,AT4G23940.1 \| FtsH extracellular protease family | 9.798896331 |

**Supplemental Table 5**. List of the unigenes that present a greater increase in their expression profiles in the vanilla root, at 10 dpi, in response to the *Fusarium* infection.

6) List top 100 of the repressed transcripts (down-regulated) at 10 dpi, in the vanilla transcriptome in response to *Fusarium*.

| **Name** | **LogFC** |
|---|---|
| TRINITY_DN82743_c0_g1_i1\|CDS1,AT3G14640.1,"AT3G14640.1 \| CYP72A10 \| cytochrome P450, family 72, subfamily A, polypeptide 10 | -10.7284808 |
| TRINITY_DN56964_c0_g1_i8\|CDS1,AT1G80840.1,"AT1G80840.1 \| WRKY40, ATWRKY40 \| WRKY DNA-binding protein 40 | -10.2747573 |
| TRINITY_DN56964_c0_g1_i8\|CDS1,AT1G80840.1,"AT1G80840.1 \| WRKY40, ATWRKY40 \| WRKY DNA-binding protein 40 | -10.2747573 |
| TRINITY_DN66527_c0_g1_i6\|CDS1,AT4G00300.2,AT4G00300.2 \| fringe-related protein | -10.1674737 |
| TRINITY_DN56964_c0_g1_i10\|CDS1,AT1G80840.1,"AT1G80840.1 \| WRKY40, ATWRKY40 \| WRKY DNA-binding protein 40 " | -9.99818758 |
| TRINITY_DN56964_c0_g1_i10\|CDS1,AT1G80840.1,"AT1G80840.1 \| WRKY40, ATWRKY40 \| WRKY DNA-binding protein 40 | -9.99818758 |
| TRINITY_DN65176_c0_g1_i3\|CDS1,AT3G01540.4,"AT3G01540.4 \| DRH1, ATDRH1 \| DEAD box RNA helicase 1 | -9.95357573 |
| TRINITY_DN65461_c0_g1_i5\|CDS1,AT4G35030.3,AT4G35030.3 \| Protein kinase superfamily protein | -9.87676324 |
| TRINITY_DN54000_c0_g1_i1\|CDS1,AT5G62020.1,"AT5G62020.1 \| AT-HSFB2A, HSFB2A \| heat shock transcription factor B2A | -9.24027043 |
| TRINITY_DN54000_c0_g1_i1\|CDS1,AT5G62020.1,"AT5G62020.1 \| AT-HSFB2A, HSFB2A \| heat shock transcription factor B2A | -9.24027043 |
| TRINITY_DN53274_c0_g1_i1\|CDS1,AT2G31390.1,AT2G31390.1 \| pfkB-like carbohydrate kinase family protein | -9.01419407 |
| TRINITY_DN53274_c0_g1_i1\|CDS1,AT2G31390.1,AT2G31390.1 \| pfkB-like carbohydrate kinase family protein | -9.01419407 |
| TRINITY_DN65011_c1_g1_i2\|CDS1,AT2G30780.1,AT2G30780.1 \| Tetratricopeptide repeat (TPR)-like superfamily protein | -9.00632806 |
| TRINITY_DN49737_c0_g1_i2\|CDS1,AT5G56260.1,AT5G56260.1 \| Ribonuclease E inhibitor RraA/Dimethylmenaquinone methyltransferase | -9.00074927 |
| TRINITY_DN49737_c0_g1_i2\|CDS1,AT5G56260.1,AT5G56260.1 \| Ribonuclease E inhibitor RraA/Dimethylmenaquinone methyltransferase | -9.00074927 |

| | |
|---|---|
| TRINITY_DN66491_c0_g1_i7\|CDS1,AT1G11330.1,AT1G11330.1 \| S-locus lectin protein kinase family protein | -8.94015998 |
| TRINITY_DN64810_c0_g1_i1\|CDS1,AT2G38110.1,"AT2G38110.1 \| ATGPAT6, GPAT6 \| glycerol-3-phosphate acyltransferase 6 | -8.93663628 |
| TRINITY_DN59211_c0_g1_i3\|CDS1,AT1G56500.1,AT1G56500.1 \| haloacid dehalogenase-like hydrolase family protein | -8.89481206 |
| TRINITY_DN59211_c0_g1_i3\|CDS1,AT1G56500.1,AT1G56500.1 \| haloacid dehalogenase-like hydrolase family protein | -8.89481206 |
| TRINITY_DN41384_c0_g1_i1\|CDS1 AT3G02740.1 AT3G02740.1\|Eukaryoticaspartylproteasefamilyprotein | -8.85773963 |
| TRINITY_DN41384_c0_g1_i1\|CDS1 AT3G02740.1 AT3G02740.1\|Eukaryoticaspartylproteasefamilyprotein | -8.85773963 |
| TRINITY_DN91391_c0_g1_i1\|CDS1,AT3G17770.1,AT3G17770.1 \| Dihydroxyacetone kinase | -8.77310227 |
| TRINITY_DN105058_c0_g1_i1\|CDS1 AT2G29940.1 AT2G29940.1\|PDR3,ATPDR3\|pleiotropicdrugresistance3 | -8.7479076 |
| TRINITY_DN105058_c0_g1_i1\|CDS1 AT2G29940.1 AT2G29940.1\|PDR3,ATPDR3\|pleiotropicdrugresistance3 | -8.7479076 |
| TRINITY_DN48321_c0_g1_i1\|CDS1,AT4G33680.1,AT4G33680.1 \| AGD2 \| Pyridoxal phosphate (PLP)-dependent transferases superfamily protein | -8.70914519 |
| TRINITY_DN48321_c0_g1_i1\|CDS1,AT4G33680.1,AT4G33680.1 \| AGD2 \| Pyridoxal phosphate (PLP)-dependent transferases superfamily protein | -8.70914519 |
| TRINITY_DN65617_c0_g1_i6\|CDS1,AT2G13440.1,AT2G13440.1 \| glucose-inhibited division family A protein | -8.6620802 |
| TRINITY_DN66144_c0_g1_i3\|CDS1,AT4G24560.1,AT4G24560.1 \| UBP16 \| ubiquitin-specific protease 16 | -8.65399793 |
| TRINITY_DN65779_c0_g1_i7\|CDS1,AT4G21300.1,AT4G21300.1 \| Tetratricopeptide repeat (TPR)-like superfamily protein | -8.63292039 |
| TRINITY_DN54683_c0_g1_i1\|CDS1,AT5G66430.1,AT5G66430.1 \| S-adenosyl-L-methionine-dependent methyltransferases superfamily protein | -8.60536746 |
| TRINITY_DN54683_c0_g1_i1\|CDS1,AT5G66430.1,AT5G66430.1 \| S-adenosyl-L-methionine-dependent methyltransferases superfamily protein | -8.60536746 |
| TRINITY_DN55626_c0_g1_i2\|CDS1,AT2G46400.1,"AT2G46400.1 \| WRKY46, ATWRKY46 \| WRKY DNA-binding protein 46 | -8.56786622 |
| TRINITY_DN55626_c0_g1_i2\|CDS1,AT2G46400.1,"AT2G46400.1 \| WRKY46, ATWRKY46 \| WRKY DNA-binding protein 46 | -8.56786622 |
| TRINITY_DN64609_c0_g1_i2\|CDS1,AT2G31955.2,AT2G31955.2 \| CNX2 \| cofactor of nitrate reductase and xanthine dehydrogenase 2 | -8.44306171 |
| TRINITY_DN66456_c0_g1_i1\|CDS1,AT2G34780.1,"AT2G34780.1 \| EMB1611, MEE22 \| maternal effect embryo arrest 22 | -8.42134686 |
| TRINITY_DN56964_c0_g1_i15\|CDS1,AT1G80840.1,"AT1G80840.1 \| WRKY40, ATWRKY40 \| WRKY DNA-binding protein 40 | -8.4078462 |

| | |
|---|---|
| TRINITY_DN56964_c0_g1_i15\|CDS1,AT1G80840.1,"AT1G80840.1 \| WRKY40, ATWRKY40 \| WRKY DNA-binding protein 40 | -8.4078462 |
| TRINITY_DN55626_c0_g1_i4\|CDS1,AT2G46400.1,"AT2G46400.1 \| WRKY46, ATWRKY46 \| WRKY DNA-binding protein 46 | -8.38864797 |
| TRINITY_DN55626_c0_g1_i4\|CDS1,AT2G46400.1,"AT2G46400.1 \| WRKY46, ATWRKY46 \| WRKY DNA-binding protein 46 | -8.38864797 |
| TRINITY_DN30000_c0_g2_i1\|CDS1 AT4G17030.1 AT4G17030.1\|ATEXLB1,EXPR,AT-EXPR,ATEXPR1,ATHEXPBETA3.1,EXLB1\|expansin-likeB1 | -8.38659777 |
| TRINITY_DN30000_c0_g2_i1\|CDS1 AT4G17030.1 AT4G17030.1\|ATEXLB1,EXPR,AT-EXPR,ATEXPR1,ATHEXPBETA3.1,EXLB1\|expansin-likeB1 | -8.38659777 |
| TRINITY_DN63645_c0_g1_i3\|CDS1,AT3G08650.1,AT3G08650.1 \| ZIP metal ion transporter family | -8.35841414 |
| TRINITY_DN63102_c0_g1_i3\|CDS1,AT5G05570.2,AT5G05570.2 \| transducin family protein / WD-40 repeat family protein | -8.33111377 |
| TRINITY_DN57080_c0_g1_i1\|CDS1,AT2G27110.3,AT2G27110.3 \| FRS3 \| FAR1-related sequence 3 | -8.31231474 |
| TRINITY_DN57080_c0_g1_i1\|CDS1,AT2G27110.3,AT2G27110.3 \| FRS3 \| FAR1-related sequence 3 | -8.31231474 |
| TRINITY_DN66141_c0_g1_i2\|CDS1,AT3G33530.1,AT3G33530.1 \| Transducin family protein / WD-40 repeat family protein | -8.26796891 |
| TRINITY_DN65967_c0_g1_i3\|CDS1,AT2G20050.1,AT2G20050.1 \| protein serine/threonine phosphatases;protein kinases;catalytics;cAMP-dependent protein kinase regulators;ATP binding;protein serine/threonine phosphatases | -8.25534136 |
| TRINITY_DN42431_c0_g1_i2\|CDS1 AT4G15630.1 AT4G15630.1\|Uncharacterisedproteinfamily(UPF0497) | -8.21728581 |
| TRINITY_DN42431_c0_g1_i2\|CDS1 AT4G15630.1 AT4G15630.1\|Uncharacterisedproteinfamily(UPF0497) | -8.21728581 |
| TRINITY_DN65890_c0_g1_i7\|CDS1,AT3G45830.1,AT3G45830.1 \| unknown protein | -8.16046939 |
| TRINITY_DN56624_c0_g1_i5\|CDS1,AT1G33490.1,AT1G33490.1 \| unknown protein | -8.15001665 |
| TRINITY_DN56624_c0_g1_i5\|CDS1,AT1G33490.1,AT1G33490.1 \| unknown protein | -8.15001665 |
| TRINITY_DN65465_c0_g1_i5\|CDS1,AT5G58040.1,"AT5G58040.1 \| ATFIP1[V], FIPS5, ATFIPS5, FIP1[V] \| homolog of yeast FIP1 [V] \| | -8.14426464 |
| TRINITY_DN58630_c0_g1_i17\|CDS1,AT3G18670.1,AT3G18670.1 \| Ankyrin repeat family protein | -8.11132905 |
| TRINITY_DN58630_c0_g1_i17\|CDS1,AT3G18670.1,AT3G18670.1 \| Ankyrin repeat family protein | -8.11132905 |
| TRINITY_DN66524_c0_g1_i5\|CDS1,AT4G35780.1,AT4G35780.1 \| ACT-like protein tyrosine kinase family protein | -8.10641732 |
| TRINITY_DN64724_c0_g1_i10\|CDS1,AT4G35600.2,AT4G35600.2 \| CONNEXIN 32 \| Protein kinase superfamily protein | -8.09121432 |
| TRINITY_DN64191_c0_g2_i3\|CDS1,AT1G24764.1,"AT1G24764.1 \| ATMAP70-2, MAP70-2 \| microtubule-associated proteins 70-2 | -8.00200031 |
| TRINITY_DN54699_c0_g1_i4\|CDS1,AT5G26667.3,AT5G26667.3 \| PYR6 \| P-loop containing nucleoside triphosphate hydrolases superfamily protein | -7.97829614 |

| | |
|---|---|
| TRINITY_DN54699_c0_g1_i4\|CDS1,AT5G26667.3,AT5G26667.3 \| PYR6 \| P-loop containing nucleoside triphosphate hydrolases superfamily protein | -7.97829614 |
| TRINITY_DN65358_c0_g1_i1\|CDS1,AT3G15920.1,AT3G15920.1 \| Phox (PX) domain-containing protein | -7.96264614 |
| TRINITY_DN65394_c0_g1_i5\|CDS1,AT1G73805.1,AT1G73805.1 \| Calmodulin binding protein-like | -7.96143327 |
| TRINITY_DN63311_c0_g1_i3\|CDS1,AT1G60060.1,AT1G60060.1 \| Serine/threonine-protein kinase WNK (With No Lysine)-related | -7.91668172 |
| TRINITY_DN58576_c0_g1_i1\|CDS1,AT5G59740.1,AT5G59740.1 \| UDP-N-acetylglucosamine (UAA) transporter family | -7.90454897 |
| TRINITY_DN58576_c0_g1_i1\|CDS1,AT5G59740.1,AT5G59740.1 \| UDP-N-acetylglucosamine (UAA) transporter family | -7.90454897 |
| TRINITY_DN40911_c0_g1_i1\|CDS1 AT3G13540.1 AT3G13540.1\|ATMYB5,MYB5\|mybdomainprotein5 | -7.90058718 |
| TRINITY_DN40911_c0_g1_i1\|CDS1 AT3G13540.1 AT3G13540.1\|ATMYB5,MYB5\|mybdomainprotein5 | -7.90058718 |
| TRINITY_DN66168_c0_g1_i9\|CDS1,AT4G01800.1,"AT4G01800.1 \| AGY1, AtcpSecA, SECA1 \| Albino or Glassy Yellow 1 | -7.89885414 |
| TRINITY_DN45953_c0_g1_i4\|CDS1 AT1G27620.1 AT1G27620.1\|HXXXD-typeacyl-transferasefamilyprotein | -7.87206362 |
| TRINITY_DN45953_c0_g1_i4\|CDS1 AT1G27620.1 AT1G27620.1\|HXXXD-typeacyl-transferasefamilyprotein | -7.87206362 |
| TRINITY_DN56964_c0_g1_i4\|CDS1,AT1G80840.1,"AT1G80840.1 \| WRKY40, ATWRKY40 \| WRKY DNA-binding protein 40 | -7.83593456 |
| TRINITY_DN56964_c0_g1_i4\|CDS1,AT1G80840.1,"AT1G80840.1 \| WRKY40, ATWRKY40 \| WRKY DNA-binding protein 40 | -7.83593456 |
| TRINITY_DN56964_c0_g1_i13\|CDS1,AT1G80840.1,"AT1G80840.1 \| WRKY40, ATWRKY40 \| WRKY DNA-binding protein 40 | -7.82036306 |
| TRINITY_DN56964_c0_g1_i13\|CDS1,AT1G80840.1,"AT1G80840.1 \| WRKY40, ATWRKY40 \| WRKY DNA-binding protein 40 | -7.82036306 |
| TRINITY_DN64550_c0_g1_i5\|CDS1,AT2G19430.1,"AT2G19430.1 \| DWA1, THO6, AtTHO6 \| DWD (DDB1-binding WD40 protein) hypersensitive to ABA 1 | -7.68770194 |
| TRINITY_DN26635_c0_g1_i1\|CDS1 AT1G64780.1 AT1G64780.1\|ATAMT1;2,AMT1;2\|ammoniumtransporter1;2 | -7.64490466 |
| TRINITY_DN26635_c0_g1_i1\|CDS1 AT1G64780.1 AT1G64780.1\|ATAMT1;2,AMT1;2\|ammoniumtransporter1;2 | -7.64490466 |
| TRINITY_DN65550_c0_g1_i2\|CDS1,AT1G66120.1,AT1G66120.1 \| AMP-dependent synthetase and ligase family protein | -7.59727774 |
| TRINITY_DN65373_c0_g1_i6\|CDS1,AT1G21730.1,AT1G21730.1 \| P-loop containing nucleoside triphosphate hydrolases superfamily protein | -7.56039678 |
| TRINITY_DN63990_c0_g1_i2\|CDS1,AT1G67850.2,AT1G67850.2 \| Protein of unknown function (DUF707) | -7.51463144 |
| TRINITY_DN46734_c0_g1_i4\|CDS1 AT1G01490.2 AT1G01490.2\|Heavymetaltransport/detoxificationsuperfamilyprotein | -7.48593857 |

| | |
|---|---|
| TRINITY_DN46734_c0_g1_i4\|CDS1 AT1G01490.2 AT1G01490.2\|Heavymetaltransport/detoxificationsuperfamilyprotein | -7.48593857 |
| TRINITY_DN64027_c0_g1_i3\|CDS1,AT3G17740.1,AT3G17740.1 \| unknown protein | -7.4384933 |
| TRINITY_DN45203_c0_g1_i1\|CDS1 AT5G19630.1 AT5G19630.1\|alpha/beta-Hydrolasessuperfamilyprotein | -7.4015463 |
| TRINITY_DN45203_c0_g1_i1\|CDS1 AT5G19630.1 AT5G19630.1\|alpha/beta-Hydrolasessuperfamilyprotein | -7.4015463 |
| TRINITY_DN65617_c0_g1_i1\|CDS1,AT2G13440.1,AT2G13440.1 \| glucose-inhibited division family A protein | -7.35746054 |
| TRINITY_DN63249_c0_g1_i2\|CDS1,AT1G35510.1,AT1G35510.1 \| O-fucosyltransferase family protein | -7.34335261 |
| TRINITY_DN66333_c0_g1_i7\|CDS1,AT1G79740.1,AT1G79740.1 \| hAT transposon superfamily | -7.32505359 |
| TRINITY_DN66417_c0_g1_i1\|CDS1,AT1G04390.1,AT1G04390.1 \| BTB/POZ domain-containing protein | -7.30241336 |
| TRINITY_DN58461_c0_g2_i4\|CDS1,AT3G63140.1,AT3G63140.1 \| CSP41A \| chloroplast stem-loop binding protein of 41 kDa | -7.29698342 |
| TRINITY_DN58461_c0_g2_i4\|CDS1,AT3G63140.1,AT3G63140.1 \| CSP41A \| chloroplast stem-loop binding protein of 41 kDa | -7.29698342 |
| TRINITY_DN63730_c0_g1_i2\|CDS1,AT5G16340.1,AT5G16340.1 \| AMP-dependent synthetase and ligase family protein | -7.21647441 |
| TRINITY_DN65096_c0_g1_i14\|CDS1,AT3G18380.2,AT3G18380.2 \| sequence-specific DNA binding transcription factors;sequence-specific DNA binding | -7.21550654 |
| TRINITY_DN53729_c0_g1_i4\|CDS1,AT1G80840.1,"AT1G80840.1 \| WRKY40, ATWRKY40 \| WRKY DNA-binding protein 40 | -7.05176914 |
| TRINITY_DN53729_c0_g1_i4\|CDS1,AT1G80840.1,"AT1G80840.1 \| WRKY40, ATWRKY40 \| WRKY DNA-binding protein 40 | -7.05176914 |
| TRINITY_DN54687_c0_g1_i8\|CDS1,AT1G08230.2,AT1G08230.2 \| Transmembrane amino acid transporter family protein | -7.03679984 |
| TRINITY_DN54687_c0_g1_i8\|CDS1,AT1G08230.2,AT1G08230.2 \| Transmembrane amino acid transporter family protein | -7.03679984 |
| TRINITY_DN54000_c0_g1_i2\|CDS1,AT3G24520.1,"AT3G24520.1 \| AT-HSFC1, HSFC1 \| heat shock transcription factor C1 | -7.03616369 |
| TRINITY_DN54000_c0_g1_i2\|CDS1,AT3G24520.1,"AT3G24520.1 \| AT-HSFC1, HSFC1 \| heat shock transcription factor C1 | -7.03616369 |

**Supplemental Table 6**. List of the 100 unigenes that present a greater decrease in their expression profiles in the vanilla root, at 10 dpi, in response to the *Fusarium* infection.
